# Metabolic specialization structures gut bacterial niches and drives colorectal cancer progression

**DOI:** 10.64898/2026.03.21.712571

**Authors:** Lin-Lin Xu, Bastian Seelbinder, Zhengyuan Zhou, Ting-Hao Kuo, Tongta Sae-Ong, Stephanie Treibmann, Victoria Damerell, Alexander Brobeil, Katharina Maria Richter, Michael Müller, Adetunji T Toriola, David Shibata, Christopher I Li, Doratha A Byrd, Jane C Figueiredo, Sheetal Hardikar, Christina E. Zielinski, Annalen Bleckmann, Yueqiong Ni, Clara Correia-Melo, Michael Zimmermann, Cornelia M Ulrich, Biljana Gigic, Gianni Panagiotou

## Abstract

Despite the established association between the gut microbiome and colorectal cancer (CRC), the functional distinction between microbial passengers and drivers of CRC progression remains unresolved. Here, we collected stool, blood, as well as paired tumor, and normal mucosa tissues from seventy-seven CRC patients to characterize the systemic and localized impact of the gut microbiome on early- and late-stage CRC. By deep shotgun metagenomic sequencing, we identified distinct bacterial species and functions residing in tumor versus normal mucosa, highlighting an enrichment of oral-associated bacteria in tumor tissues. Several of these species remained undetected in the stool microbiome analysis. We further combined bacterial culturing with untargeted metabolomics of bacteria enriched in tumor and normal mucosa tissues, revealing distinct clusters of metabolic potential. Functional testing of multiple members from one cluster comprising both tumor- and mucosa-enriched species revealed *Leptotrichia wadei* as a pro-tumorigenic bacterium in a murine CRC model. Single-nucleus RNA sequencing and *in vitro* experiments further demonstrated that *L. wadei* and its secretome induces M2 macrophage polarization to promote tumor growth. Overall, our study shows that metabolic specialization structures microbial colonization niches, while species-specific metabolic outputs identify functional drivers of CRC progression, and uncovers *L. wadei* as an oncogenic bacterium in CRC.

## INTRODUCTION

The microbiome has been shown to resident in tumor tissues in colorectal cancer (CRC) and influence various aspects of CRC, including malignant transformation, tumor progression, and responses to anticancer therapies such as immunotherapy^1–3^. Mechanistic studies have focused on tumor enriched bacterial species and their causal roles in CRC tumorigenesis. *Fusobacterium nucleatum* promotes CRC by adhering to and invading epithelial cells, secreting oncogenic metabolites and inducing an immunosuppressive tumor microenvironment^4–6^. Likewise, colibactin-producing *Escherichia coli* and enterotoxigenic *Bacteroides fragilis* promote CRC via the genotoxins colibactin (clbB) and *B. fragilis* toxin (bft), respectively^7^. However, despite these advances, a comprehensive characterization of tumor-associated bacteria remains lacking, limiting our ability to distinguish functional ‘drivers’ of CRC progression from microbial ‘passengers’^8^. Addressing this challenge requires systematic, functionally informed profiling of the tumor microbiome beyond taxonomic enrichment alone^9^.

Most human studies to date have characterized the gut microbiome in CRC using stool samples, revealing associations with disease initiation, progression and treatment response^10–13^. These analyses have identified species enriched in CRC, including *F. nucleatum*, *B. fragilis* and *Parvimonas micra*^14–16^, whereas putatively protective taxa such as *Clostridium butyricum* and *Lactobacillus gallinarum* are diminished^14^. In parallel, gut microbes produce metabolites that enter the circulation and modulate tumor microenvironment and host immune responses^17–19^. Microbial metabolites, including short-chain fatty acids, formate, and branched-chain amino acids, have been implicated in immune cell activation, inflammation, and tumor cell proliferation^5,19–22^. However, stool-based surveys incompletely capture tissue-resident microbes and typically lack functional resolution to link specific species to systemic metabolic and immune consequences. Although individual CRC-associated taxa have been reported, an integrated framework combining tissue microbiomes with species-level metabolic outputs and host metabolomics is still missing.

Here, we applied an integrated multi-omics approach in a cohort of 77 patients with stage I-IV CRC, profiling both stool and paired tumor and distant normal mucosa microbiomes with a focus on microbial functional potential. We integrated these microbiome data with host-and bacterial metabolomics, lipidomics and circulating cytokine markers to define metabolic signatures linked to tissue colonization niches and systemic host effects. *In vitro* culture of selected isolates enabled reconstruction of a community metabolic map for tumor- and normal mucosa-enriched microbes. Using CRC cell lines and *in vivo* mouse models, we demonstrate that species-specific metabolic outputs, rather than global community metabolic features, distinguish microbial passenger bacteria from candidate drivers of CRC progression. Specifically, we identify the oral bacterium *Leptotrichia wadei* as a candidate driver that promotes colorectal tumorigenesis by inducing M2 macrophage polarization.

## RESULTS

### Microbial Dynamics and Metabolic Shifts from Early- to Late-Stage Colorectal Cancer

To evaluate systemic and local changes associated with CRC progression, we selected patients diagnosed with primary invasive CRC from the ColoCare Study^23^, recruited at the Heidelberg University Hospital (Heidelberg, Germany). The study cohort included 77 participants (49 male, 28 female) aged from 38 to 87 years, with tumors located in either the colon (n = 29) or rectum (n = 48) and distributed across four cancer stages (Stage I: 11; Stage II: 19; Stage III: 28; Stage IV: 19) (Figure 1A; Table S1). From these 77 patients, we collected paired tumor (n = 67) and normal mucosa tissue (n = 69), as well as stool (n = 76) and serum samples (n = 77), to enable a multi-omics characterization of CRC patients.

**FIGURE 1.**
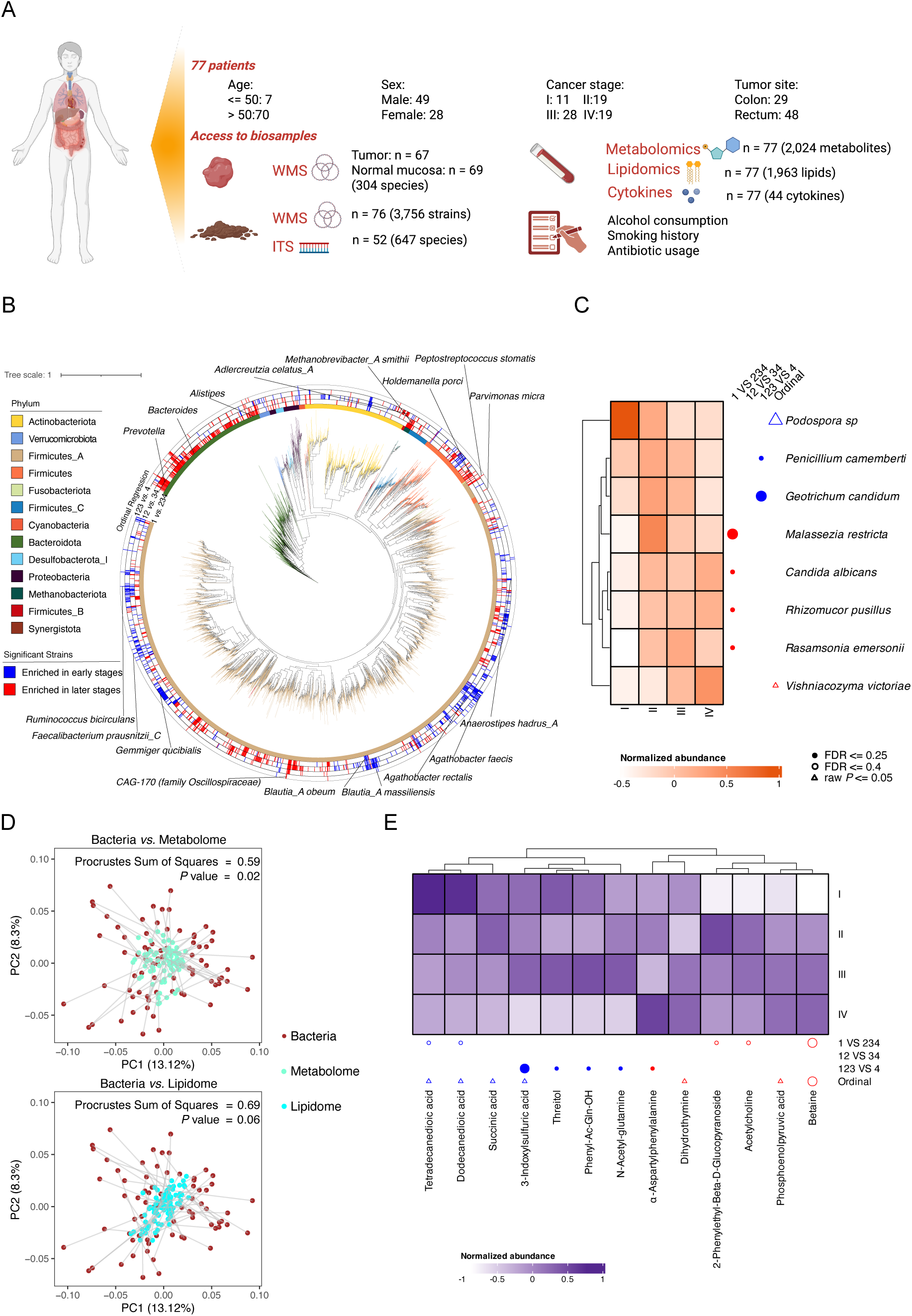
Microbial dynamics and metabolic shifts from early- to late-stage colorectal cancer. (A) Study design. Stool, paired tumor and normal mucosa tissue, and blood samples were retrieved from 77 patients across four cancer stages. Created with Biorender.com. (B) In total, 1288 differentially abundant strains are shown in a phylogenetic tree, grouped in the phyla *Firmicutes*, *Bacteroidota*, *Actinobacteriota*, *Proteobacteria* and *Fusobacteriota*. In the outer circles, strains are marked for significant elevation or depletion. Several stage-specific and stage-consistent strains are labelled. Pairwise comparisons were conducted to evaluate the stage-specific signatures in progression of colorectal cancer (CRC) across consecutive stages (I vs. II-IV, I-II vs. III-IV, and I-III vs. IV). Ordinal regression was performed to identify stage-consistent signatures showing progressive increases or decreases along CRC stages I to IV. (C) Clustered heatmap showing significantly differentially abundant fungal species across cancer progression stages. Pairwise comparisons were performed using DESeq2 (P ≤ 0.05, FDR ≤ 0.4) (I vs. II-IV, I-II vs. III-IV, and I-III vs. IV). Species consistently differing across stages were identified using ordinal regression (P ≤ 0.05). Heatmap intensities (red color scale) represent scaled and centered relative abundances. (D) Procrustes analysis showing overall association between variation in stool bacterial community and serum metabolome (upper panel) and lipidome (bottom panel). (E) Clustered heatmap showing significantly differentially abundant metabolites across cancer progression stages. Pairwise comparisons were performed using POMA (P ≤ 0.05, FDR ≤ 0.4) (I vs. II-IV, I-II vs. III-IV, and I-III vs. IV). Metabolites consistently differing across stages were identified using ordinal regression (P ≤ 0.05). Heatmap intensities (purple color scale) represent scaled and centered value of the logarithm transformed abundance.

We first performed shotgun whole-genome metagenomics on stool samples to systematically characterize the gut microbiome of CRC patients. Alpha and beta diversity showed no significant differences across the four cancer stages (Wilcoxon rank-sum test, P > 0.05; PERMANOVA, P > 0.05) (Figure S1A and S1B). Smoking and antibiotic usage were significantly influencing stool bacterial community composition (PERMANOVA, P ≤ 0.05) (Figure S1C). We next constructed dereplicated metagenome-assembled genomes (MAGs) of the stool microbiota, identifying 3756 MAGs, and used them to profile strain-level shifts between early and late CRC (DESeq2, FDR ≤ 0.05) (Figure 1B). In total, 494 MAGs were enriched in early CRC and 706 in late CRC (88 showed mixed enrichment across contrasts). Comparing our results to the most comprehensive metagenomic meta-analysis of CRC to date^13^, 280 of our stage-associated MAGs mapped to 52 species-level genome bins (SGBs) and showed consistent abundance patterns. In particular, *Peptostreptococcus stomatis*, *P. micra* and *Methanobrevibacter smithii* have been reported as CRC-enriched taxa^13^, and were relatively more abundant in late-stage CRC in our cohort (DESeq2, FDR ≤ 0.05). By contrast, *Faecalibacterium prausnitzii*, *Anaerostipes hadrus*, *Agathobacter faecis* and *Ruminococcus bicirculans* have been reported as enriched in healthy controls^13^, and were relatively more abundant in early-stage CRC in our cohort (DESeq2, FDR ≤ 0.05).

Beyond the shared signatures, a large number of *Bacteroidota* strains (n = 231) were significantly enriched in late-stage cancer of our cohort (stages II–IV; DESeq2, FDR ≤ 0.05), primarily from the genera *Prevotella* (*Prevotella copri*), *Bacteroides* (*B. fragilis*) and *Alistipes* (*Alistipes onderdonkii*). Additional strains enriched in late-stage CRC included *Holdemanella porci* and *CAG-170* (family *Oscillospiraceae*) (stages II to IV; DESeq2, FDR ≤ 0.05). *Adlercreutzia celatus_A* were enriched exclusively in stage IV (DESeq2, FDR ≤ 0.05). In contrast, *Agathobacter rectalis* decreased gradually from stage I to IV (ordinal regression, P ≤ 0.05). Additional strains enriched in early-stage cancer included *Blautia_A massiliensis* and *Blautia_A obeum* (stages I; DESeq2, FDR ≤ 0.05), as well as *Gemmiger qucibialis*, which was depleted exclusively in stage IV (DESeq2, FDR ≤ 0.05).

Given that fungal contributions to CRC are largely unexplored, we next performed full-length internal transcribed spacer (ITS) sequencing to characterize the fungal community in 52 stool samples that passed quality control. In total, we identified 647 fungal species, which were reduced to 109 species after filtering (see Star Methods for details). Alpha diversity showed no significant differences across the four cancer stages (Wilcoxon rank-sum test, *P* > 0.05) (Figure S1A). In contrast, beta diversity analysis revealed significant differences in the fungal community composition among the four cancer stages at both the genus and species levels (PERMANOVA, *P* ≤ 0.05) (Figure S1B). Specifically, stage I differed significantly from late cancer stages (PERMANOVA, *P* ≤ 0.05). Additionally, stool fungal community composition was also shaped by antibiotic usage (PERMANOVA, *P* ≤ 0.05) (Figure S1C). This is consistent with our previous work showing that antibiotic exposure can induce a long-lasting disruption of the gut mycobiome^24^. Specific fungal species—including *Malassezia restricta*, *Candida albicans*, *Rhizomucor pusillus*, and *Rasamsonia emersonii*—were more abundant in late cancer stages compared to stage I (DESeq2, *P* ≤ 0.05, FDR ≤ 0.25), whereas *Penicillium camemberti* and *Geotrichum candidum* were decreased in stage IV (Figure 1C).

We then performed untargeted liquid chromatography–mass spectrometry (LC–MS)-based metabolomics and lipidomics on serum samples to further characterize CRC patients. This analysis revealed a significant correspondence between bacterial community variation and serum metabolites across subjects (Procrustes analysis, P ≤ 0.05; Figure 1D), but not with the serum lipid profile. We next identified serum metabolomics signatures related to cancer stages (Figure 1E). Dihydrothymine, phosphoenolpyruvic acid, and betaine increased progressively from stage I to stage IV (ordinal regression, P ≤ 0.05), whereas 3-indoxylsulfuric acid, succinic acid, tetradecanedioic acid, and dodecanedioic acid decreased across stages (ordinal regression, P ≤ 0.05). Acetylcholine was enriched in late cancer stages (stages II–IV; POMA, FDR ≤ 0.4). In addition, 3-indoxylsulfuric acid, threitol, phenyl-Ac-Gln-OH, and N-acetyl-glutamine exhibited lower abundances in stage IV, while α-aspartylphenylalanine was increased (POMA, FDR ≤ 0.25).

These analyses reveal that gut bacterial communities exhibit stage-specific shifts in CRC, which correspond closely with serum metabolite profile, highlighting a systemic microbiome–host metabolic signature, while fungal communities show stage-specific changes without direct correlation to host metabolic profiles.

### Stage-Specific Metabolic Signatures Define CRC Microbial Clusters

We subsequently performed Kyoto Encyclopedia of Genes and Genomes (KEGG) Orthology (KO)-based clustering of the early- and late-stage enriched bacterial signature strains, identifying in total ten functional clusters (Figure S2), seven of which were “pure clusters” (Figure 2A): four dominated by late-stage–enriched strains and three by early-stage–enriched strains. Cluster C2 was dominated by *F. prausnitzii* strains, C4 by *G. qucibialis*, and C6 by *B. massiliensis* and *B. obeum*. Cluster C5 was dominated by *H. porci*. Cluster C8 was dominated by *Firmicutes* strains, including *P. stomatis*, *P. micra* and *CAG-170* (family *Oscillospiraceae*), whereas C9 was dominated by *Bacteroidota* strains. Cluster C10 comprised archaeal methanogens, including *Methanobrevibacter_A smithii_A*, *M. smithii* and *Methanosphaera stadtmanae*.

**FIGURE 2.**
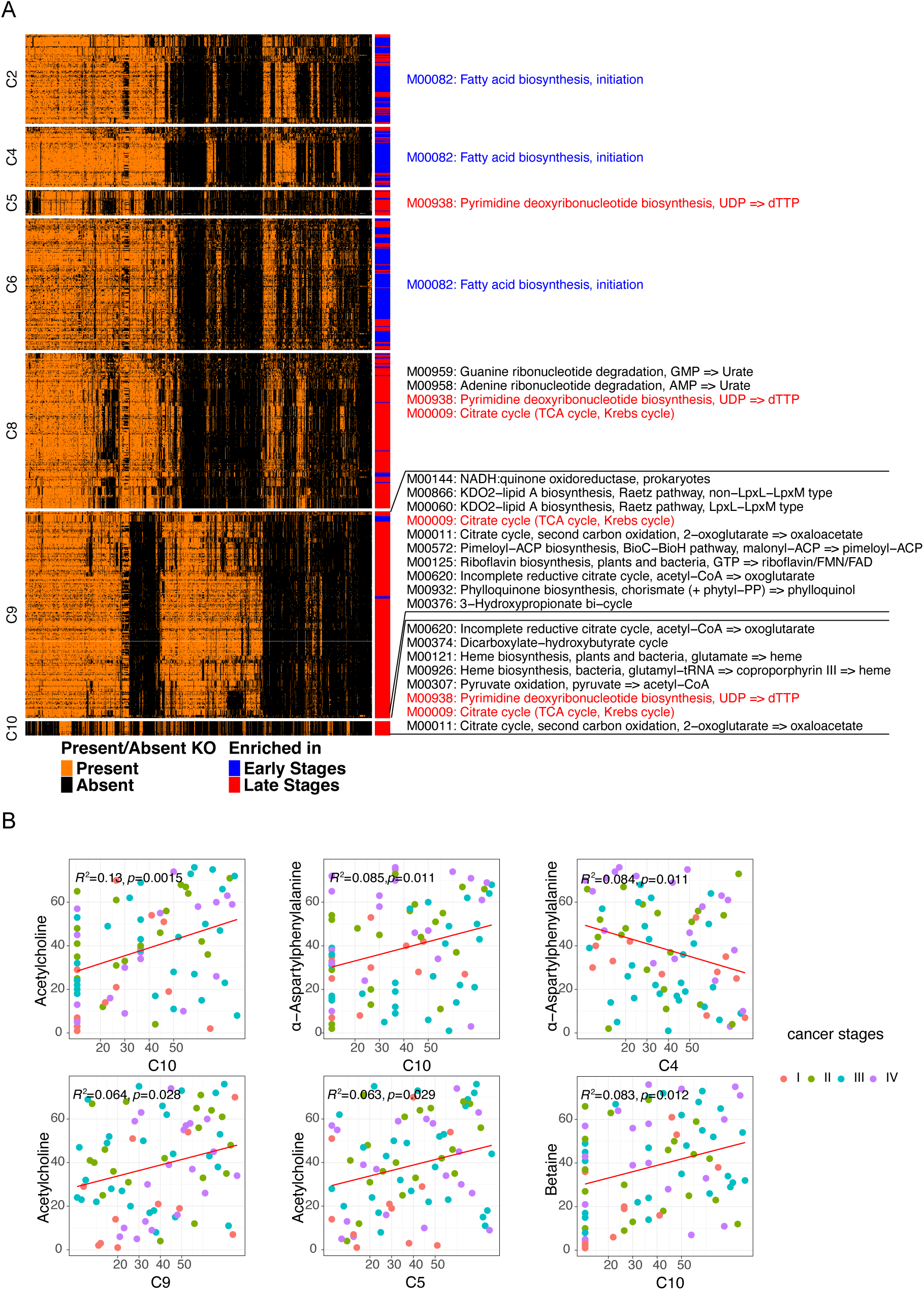
Stage-specific metabolic signatures define CRC microbial clusters. (A) Presence/absence matrix of Kyoto Encyclopedia of Genes and Genomes (KEGG) Orthologies (KOs) identified from signature bacterial strains. The strains are clustered by Partitioning Around Medoids (PAM) algorithm, and for each cluster the KEGG pathways most uniquely enriched in early- or late-stage–associated clusters are shown alongside. (B) Scatter plot demonstrating significant Spearman correlations (r) and p values between functional clusters and serum metabolites. Two-tailed Spearman’s rho correlation testing was used and linear regression lines shown.

Over-representation analysis (FDR ≤ 0.05) revealed that the fatty acid biosynthesis module (M00082) was significantly enriched in clusters dominated by early-stage–enriched strains. In contrast, late-stage–dominated clusters were associated with nucleotide biosynthesis (pyrimidine deoxyribonucleotide biosynthesis) and central carbon and energy metabolism (citrate cycle/ tricarboxylic acid (TCA) cycle). Specifically, clusters C5, C8 and C10 were significantly enriched for the pyrimidine deoxyribonucleotide biosynthesis module (M00938) (FDR ≤ 0.05), a component of the broader pyrimidine metabolism pathway that is conserved between host and gut bacteria and is essential for DNA replication and RNA synthesis, thereby supporting cancer cell proliferation^25^. Furthermore, late-stage–dominated clusters C8, C9 and C10 were significantly enriched for the citrate cycle (TCA cycle, Krebs cycle; M00009) (FDR ≤ 0.05), and cluster C10 showed a unique enrichment of the heme biosynthesis module (M00926).

To delineate the associations between bacterial function clusters and serum metabolomics signatures related to cancer stages, we applied Spearman’s rank correlation analysis (Figure 2B). Functional cluster C10 was significantly positively correlated with acetylcholine, α-aspartylphenylalanine and betaine, whereas cluster C4 was negatively correlated with α-aspartylphenylalanine. Clusters C9 and C5 showed positive correlations with acetylcholine. In contrast, clusters C2, C6, and C8, despite showing stage-specific enrichment, did not exhibit significant correlations with serum metabolites.

Our findings reveal that early- and late-stage–enriched stool bacteria and archaea resolve into functional clusters with distinct metabolic focuses that associate with specific serum metabolites linked to CRC progression, supporting these clusters as plausible candidates for systemic functional drivers.

### Oral-associated Bacteria Undetected in Stool Microbiota Characterize Functional Signatures in CRC Tumors

To explore the role of the intra-tissue microbiome in driving the physiological states of tumor tissues in CRC patients, we collected paired tumor, and normal mucosa tissues from each CRC patient for deep shotgun metagenome sequencing. Our analysis showed significant correspondence between microbiome community from tumor, normal mucosa, and stool samples (Procrustes, P ≤ 0.05) (Figure 3A). Tumor and normal mucosa microbiome communities are significant different (Figure 3B) (PERMANOVA test, P = 0.001).

**FIGURE 3.**
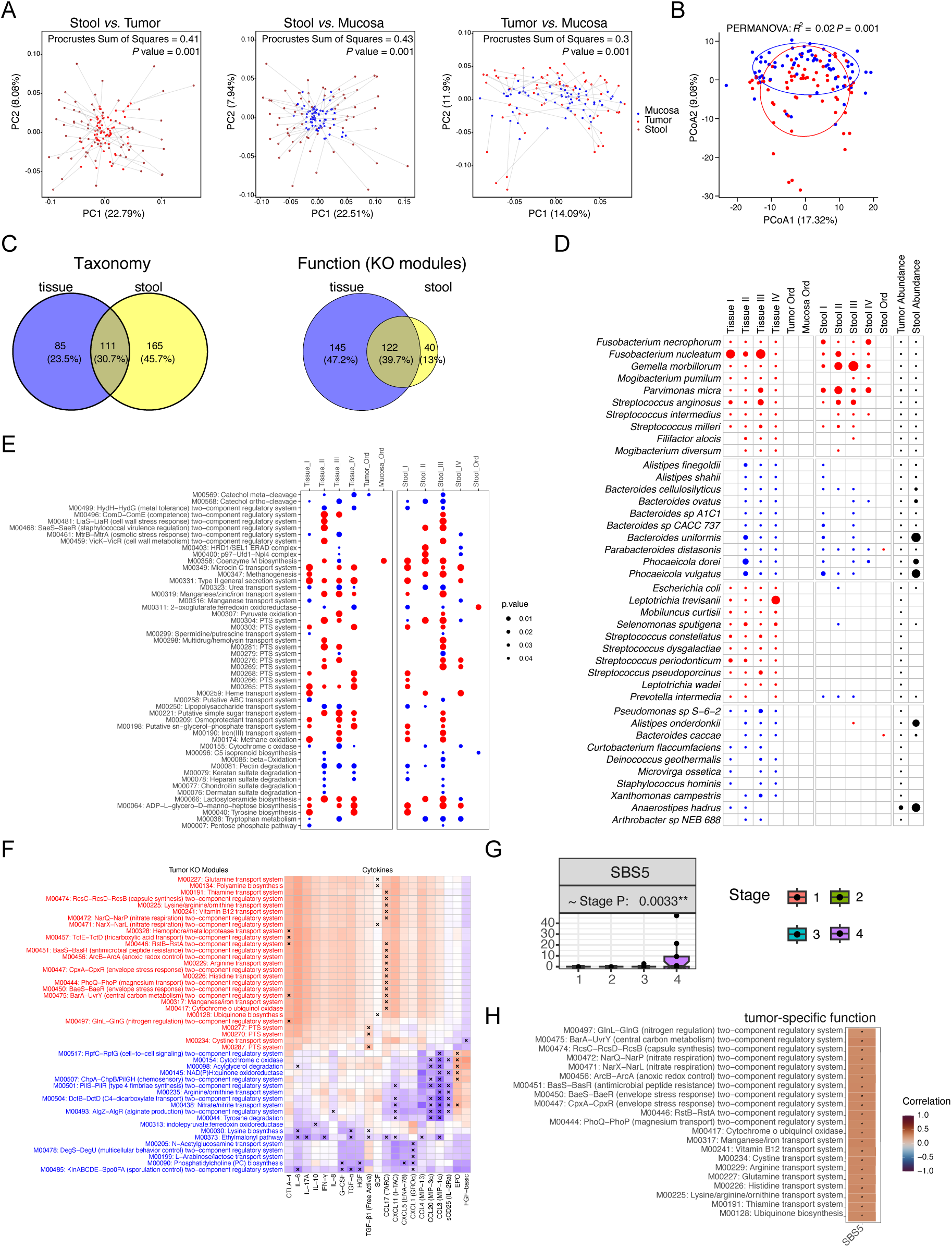
Oral-associated bacteria undetected in stool microbiota characterize functional signatures in CRC tumors. (A) Procrustes analysis showing overall association between variation in bacterial community compositions across sample types. Left: stool *vs.* tumor tissue; middle: stool *vs.* normal mucosa tissue; right: tumor tissue *vs.* normal mucosa. (B) PCoA of tumor and normal mucosa bacterial community. PCoA: Principal Coordinate Analysis. (C) Venn diagrams showing the overlap of bacterial signatures identified in tissue and stool samples. The left panel represents shared and unique bacterial species, while the right panel represents shared and unique KO modules. (D) Heatmap displaying the top ten significantly differential bacterial signatures between tumor and normal mucosa (tissue samples), and their patterns of change in stool samples from CRC patients and controls across four cancer stages. The top two panels display bacterial signatures detected in both tissue and stool samples, while the bottom two panels show signatures identified exclusively in tissue samples. Circle color indicates the direction of enrichment (tumor *vs.* normal), while circle intensity corresponds to the statistical significance (−log_10_P-value). Differential bacterial signatures were identified using DESeq2 and ordinal regression. DESeq2 results are considered significant with raw P ≤ 0.05 and FDR ≤ 0.25, while ordinal regression results are significant with raw P ≤ 0.05. (E) Heatmap displaying tissue- and stool-shared differential bacterial KO modules between tumor and normal mucosa (tissue samples) and CRC patients and controls (stool samples) across four cancer stages. Circle color indicates the direction of enrichment (tumor *vs.* normal; CRC *vs.* Control), while circle intensity reflects statistical significance (−log_10_P-value). Differential bacterial functions were identified using DESeq2 and ordinal regression. Results from DESeq2 were considered significant at raw P ≤ 0.05 and FDR ≤ 0.25, while ordinal regression results were considered significant at raw P ≤ 0.05. (F) Spearman correlation analysis between tissue-specific KO modules and serum cytokine levels. KO modules labeled in red are enriched in tumor tissue, while those in blue are enriched in normal mucosa. An “X” in a cell indicates a statistically significant correlation (P ≤ 0.05). (G) Human cancer mutational signature analysis showing an increase in single base substitution signature COSMIC SBS5 from cancer stages I to IV (ordinal regression: P ≤ 0.05). (H) Spearman correlation analysis between tissue-specific KO modules and human mutational signature SBS5. Only statistically significant correlations are shown (P ≤ 0.05).

Next, we aimed to investigate microbial species enriched in tumor tissues. To determine whether these features are tissue-specific or shared between tissue and stool samples, we integrated publicly available shotgun metagenomic sequencing datasets^15,26–29^ and matched them to our CRC patients (Figure S3). By comparing signatures from tissue and stool, we identified 85 tissue-specific species and 165 stool-specific species, with 111 species shared between both (Figure 3C). Among the shared signatures, *F. nucleatum*, *Fusobacterium necrophorum*, and *Gemella morbillorum* were consistently elevated in tumors and in CRC patient stool across Stages I-IV (Figure 3D). In contrast, *Parabacteroides distasonis*, *Bacteroides cellulosilyticus*, and *Phocaeicola vulgatus* were enriched in normal mucosa and controls across in at least three cancer stages (Figure 3D). Notably, we found that oral-associated bacteria, including species belonging to *Streptococcus*, *Prevotella*, and *Leptotrichia*, were enriched in tumor tissues and most of these species could not be detected in stool samples. Specifically, *Streptococcus constellatus*, *P. intermedia*, and *L. wadei* were consistently enriched in tumors across at least three cancer stages.

Considering the functional potential of the microbiome community, we identified 145 tissue-specific KO modules and 40 stool-specific KO modules, and 122 KO modules are shared (Figure 3C). Some of the shared modules are related to the phosphotransferase system (PTS) and are enriched in both tumor tissues and stool samples of CRC patients. KEGG module ADP-L-glycero-D-manno-heptose biosynthesis (M00064) is enriched in both tumor and stool of CRC patients, whereas tryptophan metabolism (M00038) is depleted (Figure 3E). KEGG modules related to amino acid transport, including the glutamine transport system (M00227), arginine transport system (M00229), histidine transport system (M00226), and lysine/arginine/ornithine transport system (M00225) as well as the anaerobic regulation module ArcB-ArcA (anoxic redox control) two-component system (M00456), were specifically enriched in tumor tissues and did not differ significantly in stool samples (Figure 3F). In contrast, the lipid metabolism modules, acylglycerol degradation module (M00098) and phosphatidylcholine (PC) biosynthesis (M00090), were specifically enriched in normal mucosal tissues. To investigate the impact of tumor-enriched and tumor-depleted microbial functions on the host immune response, we correlated KEGG modules uniquely differentially abundant in tissue with serum cytokine levels (Figure 3F). We found sixteen significant positive correlations with CCL17 (TARC), including arginine transport system (M00229), lysine/arginine/ornithine transport system (M00225), and ArcB-ArcA (anoxic redox control) two component regulatory system (M00456).

Next, we analyzed the human reads in tissue samples to identify mutational signatures in human cancer cells. Specifically, COSMIC SBS5 signature is increased from cancer stage I to stage IV (Ordinal regression, P ≤ 0.05) (Figure 3G). We further correlated these human cancer signatures to tissue-specific microbiome function signatures. SBS5 showed significant correlations with eleven modules related to the two-component regulatory system in tumors, including the anoxic redox control ArcB-ArcA two-component regulatory system (M00456) (Figure 3H). M00456 is enriched in facultative anaerobic bacteria, allowing them to adapt their metabolism in response to oxygen availability^30^, which may associated to oxygen-responsive regulatory pathways for tumor-enriched oral-associated bacteria to survive in the tumor microenvironment.

Overall, we identified tissue-specific microbial species and functions and their associations with host immune system and cancer signatures. Notably, oral-associated bacteria were implicated in both tumor taxa and functional modules (e.g., ArcB–ArcA (anoxic redox control; M00456)), suggesting a central role of oral-associated bacteria in the tumor microenvironment.

### Tissue Resident Bacterial Species Resolve into Distinct Metabolic Clusters

To identify candidate functional drivers of CRC, we selected 29 bacterial species enriched in tumor or normal mucosa, available in our strain collection, cultured them *in vitro*, and characterized their metabolic outputs using untargeted metabolomics analysis. After quality-control filtering, 3,502 of the 3,789 detected metabolites were retained for subsequent analysis. Principal component analysis revealed a significant separation between tumor- and normal mucosa-enriched bacterial species (PERMANOVA, P ≤ 0.05) (Figure 4A). The overall difference in data variance was driven by valine, cysteine, carnosine, alanine, and oxidized glutathione (projection analysis, P ≤ 0.05, FDR ≤ 0.2). The metabolic profiles of tumor-enriched and mucosa-enriched species clustered the strains into four metabolic clusters (Figure 4B). Cluster C1 comprised only mucosa-enriched strains, whereas cluster C3 consisted exclusively of tumor-enriched strains. Tumor-enriched oral-associated and non-oral-associated bacteria were intermixed in cluster C3. Cluster C4 contained four *Fusobacterium* subspecies, while cluster C2 was the only mixed group, harboring tumor-enriched species *L. wadei*, *G. morbillorum*, *P. micra*, and *B. fragilis*, together with mucosa-enriched species *A. hadrus* and *C. butyricum*. Notably, this mixed cluster included the tumor-promoting species *B. fragilis* and *P. micra*, and the tumor-suppressing *C. butyricum*^21,31,32^, which have been validated in mice models.

**FIGURE 4.**
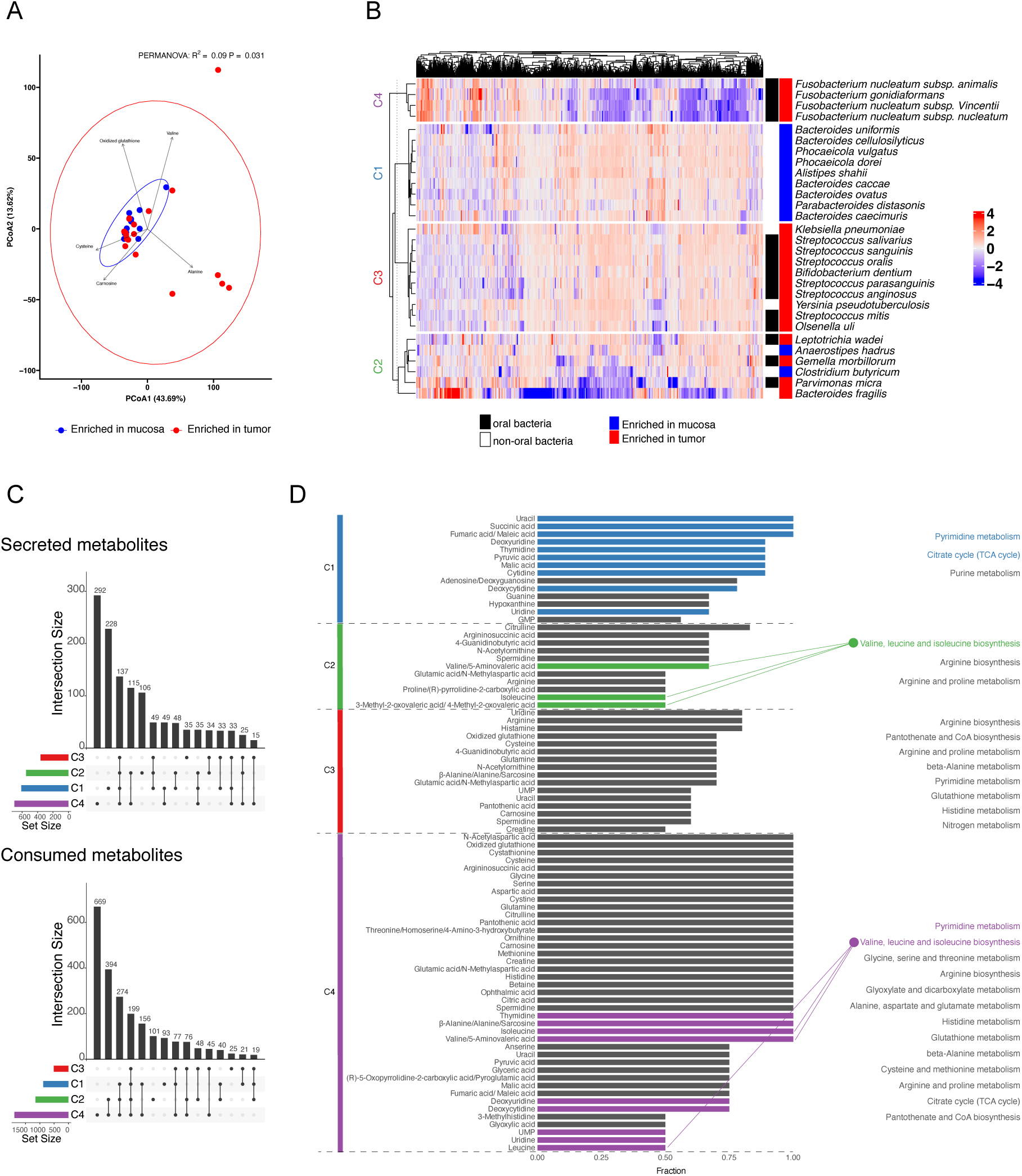
Tissue resident bacterial species resolve into distinct metabolic clusters. (A) Principal component analysis (PCA) of the metabolic profiles of tumor- and normal-mucosa-resident bacteria. (B) Normalized abundance matrix of metabolites identified from the supernatants of tumor-enriched bacteria (red) and normal mucosa-enriched bacteria (blue). Bacterial strains are grouped into four clusters using the Partitioning Around Medoids (PAM) algorithm based on their metabolic profiles. (C) UpSet plot showing the number of secreted (top panel) and consumed (bottom panel) metabolites identified via POMA (P ≤ 0.05, FDR ≤ 0.25) in each bacterial metabolic cluster and shared across combinations of clusters. The numbers above each column indicate the count of shared secreted or consumed metabolites. The set size bars on the left represent the total number of secreted or consumed metabolites in each bacterial metabolic cluster, while the connected dots below each column indicate which clusters share those metabolites. (D) Metabolic pathways enriched for secreted and consumed metabolites in each bacterial metabolic cluster (MetaboAnalyst: FDR ≤ 0.05). Red, blue, green, and purple bars represent metabolites secreted in each respective cluster, while grey bars indicate consumed metabolites. Metabolic pathways enriched by secreted metabolites are shown in the corresponding cluster colors (red, blue, green, or purple), and pathways enriched by consumed metabolites are shown in grey. Lines connecting metabolites to pathways indicate that the metabolites are components of those pathways.

We next identified metabolites that were secreted or consumed by these bacterial species (POMA, P ≤ 0.05, FDR ≤ 0.25), by comparing the metabolic profile of each species’ supernatant with its corresponding original culture medium (Figure 4C). Cluster C4 exhibited the largest set of unique secreted and consumed metabolites, underscoring the distinctive metabolic capacity of the *Fusobacterium* subspecies, whereas cluster C2 shared the greatest overlap with C4, suggesting partial convergence of metabolic functions between these two clusters. Pathway-enrichment analysis based on confidently annotated features (MetaboAnalyst, FDR ≤ 0.05; see Star Methods for details) revealed that mucosa-enriched species (C1) and *Fusobacterium* subspecies (C4) secreted metabolites enriched in pyrimidine metabolism, while tumor-enriched species (C3) consumed metabolites from pyrimidine metabolism (Figure 4D). Notably, cluster C1 showed a distinct signature of purine metabolism consumption, which was not observed in the other clusters. Species in clusters C2 and C4 secreted metabolites associated with valine, leucine, and isoleucine biosynthesis, with the mixed cluster (C2) secreting valine, isoleucine, 3-Methyl-2-oxovaleric acid (KMV) and 4-Methyl-2-oxovaleric acid (KIC), and the *Fusobacterium* cluster (C4) secreting valine, isoleucine, and leucine. In contrast, metabolites in arginine and proline metabolism pathways were consistently consumed by species in clusters C2, C3, and C4 (Figure 4D).

Collectively, these findings provided a comparative map of secreted and consumed metabolites for bacteria inhabiting colorectal tumors and normal mucosa tissues. Tissue-resident bacteria exhibit distinct metabolic clusters corresponding to tumor or mucosa enrichment, but high-level pathway enrichment shows overlap across clusters, indicating that functional drivers may be hidden in species-specific metabolic outputs. Among the clusters, the mixed cluster (C2) was the most intriguing, containing both tumor- and mucosa-enriched species including known tumor-promoting bacteria, making it a prime candidate for species-level functional validation.

### *Leptotrichia wadei* Promotes CRC Progression by Inducing M2 Macrophage Polarization

To pinpoint species-specific functional drivers within the mixed cluster (C2), which contains both tumor- and mucosa-enriched bacteria, we tested individual strains *in vitro* and *in vivo* for their ability to influence CRC growth. To test whether two mucosa-enriched species, *A. hadrus* and *C. butyricum*, affect CRC cell survival and proliferation, seven CRC cell lines as well as healthy colonic epithelial cells were treated with bacterial secretomes of these bacteria (Figure S4A). Both secretomes significantly increase the dead cell count and decrease the number of proliferating cells and the confluency increase of most analyzed CRC cell lines, especially of DLD1 and SW-480, while only having a minor effect on the healthy colonic epithelial cells. The effects of the secretomes of these two butyrate producing microbes mirror the effects of 2 mM butyrate which was previously shown to inhibit CRC cell proliferation through autophagy degradation of β-catenin^33^.

Furthermore, we investigated whether tumor-enriched oral bacteria *G. morbillorum* and *L. wadei* can promote CRC by a subcutaneous tumor model (Figure 5A). Intriguingly, injection of *L. wadei* into subcutaneous tumors significantly promoted the tumor size (Wilcoxon rank-sum test, *P* ≤ 0.05), while *G. morbillorum* showed no difference with control group (Figure 5A). While we found *L. wadei* in tumor tissue samples of our CRC patient’s, it was not detected in their stool samples, and also detected rarely (2 out of 3068 individuals; relative abundance < 0.015%) in stool samples of healthy patients^34^. To further investigate the potential mechanism, we added heat killed *L. wadei*, conditioned medium of *L. wadei*, and original culture medium groups (Figure 5B and 5C). Both alive *L. wadei* and conditioned medium of *L. wadei* significantly promoted tumor growth (Wilcoxon rank-sum test, *P* ≤ 0.05) compared to control and original culture medium group respectively, while heat killed *L. wadei* group had no significant difference with control group (Wilcoxon rank-sum test, *P* > 0.05) (Figure 5C, 5D, Figure S4B, S4C). Six *L. wadei*-injected tumors were plated, confirming the presence of viable *L. wadei* within the tumors (Table S2). Together, our results suggested that *L. wadei* secretome could promote CRC tumorigenesis.

**FIGURE 5.**
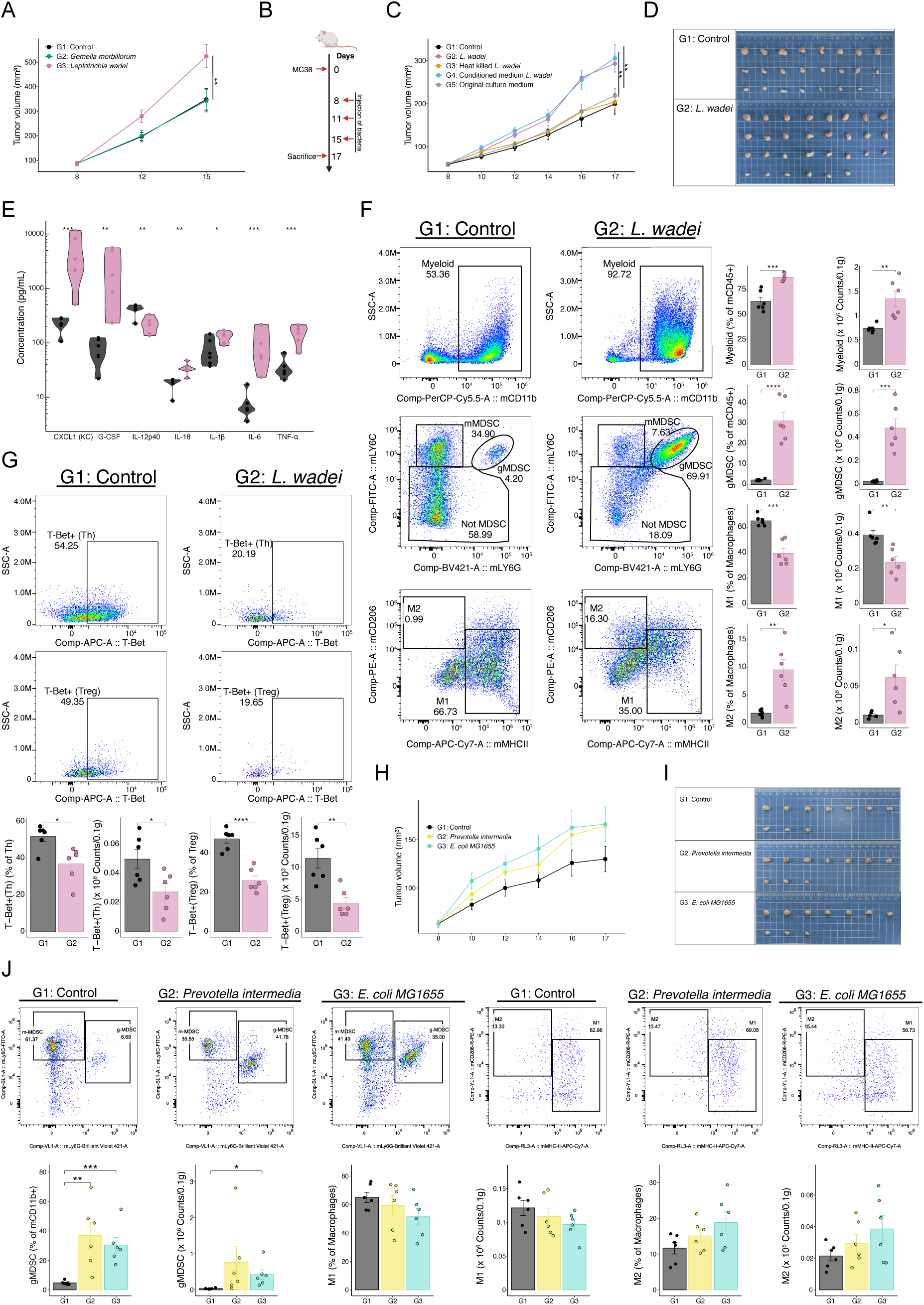
*Leptotrichia wadei* promotes CRC progression by inducing M2 macrophage polarization. (A) C57BL/6J mice were subcutaneously injected with 1.2 x 10^5^ MC38 cells. After 8 days, 1 x 10^8^ cells of the oral bacteria (*Gemella morbillorum* and *Leptotrichia wadei*) were injected into subcutaneous tumors twice a week and PBS served as a negative control. The tumor volume (mm^3^) was measured and shown as mean ± SEM. The tumor volumes of indicated groups at day 15 were compared by using Wilcoxon rank-sum test (p ≤ 0.05; *, p ≤ 0.01; **, p ≤ 0.001; ***). Each group has 8 mice. (B) Experimental design of subcutaneous tumor model. (C) C57BL/6J mice with MC38 subcutaneous tumors were treated as previously described. Mice were injected with Control (sterile PBS (with 0.5% cysteine), n = 24), *L. wadei* (1 x 10^6^ CFU) (n = 30), heat killed *L. wadei* (n = 20), *L. wadei* conditioned medium (n = 15), or original culture medium (n = 15) twice a week. The tumor volumes of indicated groups at day 17 were compared by using Wilcoxon rank-sum test. (D) Images of representative MC38 subcutaneous tumors after the intra-tumoral injection of the indicated groups in C57BL/6J mice. (E) Concentration of cytokines in tumor tissue following intra-tumoral injection. Statistical significance between the *L. wadei* and control groups was evaluated using a two-sided Student’s t-test. Each group included six mice. (F) The tumor-infiltrating myeloid cells (mCD45+mCD11b+), M1 macrophage (mCD45+mCD11b+nF4/80+MHCIIhiCD206lo), M2 macrophage (mCD45+mCD11b+nF4/80+MHCIIloCD206hi), and gMDSC cells (mCD45+mCD11b+mLy6G+mLy6Clo) in MC38 subcutaneous tumors from mice with intra-tumoral injection of indicated bacteria were analyzed by flow cytometry. The percentage and cell number were shown as mean ± SEM. Statistical significance between the *L. wadei* and control groups was evaluated using a two-sided Student’s t-test. Each group included six mice. (G) The tumor-infiltrating T-Bet+ regulatory T (Treg) cells (mCD45+mCD11b-mCD3+mCD4+mCD25+FOXP3+T-Bet+) and T-Bet+ helper T (Th) cells (mCD45+mCD11b-mCD3+mCD4+mCD8-) in MC38 subcutaneous tumors from mice with intra-tumoral injection of indicated bacteria were analyzed by flow cytometry. The percentage and cell number were shown as mean ± SEM. Statistical significance between the *L. wadei* and control groups was evaluated using a two-sided Student’s t-test. Each group included six mice. (H) C57BL/6J mice with MC38 subcutaneous tumors were treated as previously described. Mice were injected with gram negative bacteria (*Prevotella intermedia* and *E. coli MG1655*). The tumor volumes of indicated groups at day 17 were compared by using Wilcoxon rank-sum test. Each group has 10 mice. (I) Images of representative MC38 subcutaneous tumors after the intra-tumoral injection of the indicated groups in C57BL/6J mice. (J) The tumor-infiltrating gMDSC cells (mCD45+mCD11b+mLy6G+mLy6Clo), M1 macrophage (mCD45+mCD11b+nF4/80+MHCIIhiCD206lo), and M2 macrophage (mCD45+mCD11b+nF4/80+MHCIIloCD206hi) in MC38 subcutaneous tumors from mice with intra-tumoral injection of indicated bacteria were analyzed by flow cytometry. The percentage and cell number were shown as mean ± SEM. Statistical significance between the indicated groups was evaluated using a two-sided Student’s t-test. Each group included six mice.

We next explored tumor cytokines regulated by *L. wadei* (Figure 5E). *L. wadei* upregulated the pro-inflammatory cytokines chemokine (C-X-C motif) ligand 1 (CXCL1), granulocyte colony stimulating factor (G-CSF), interleukin-18 (IL-18), interleukin-1 beta (IL-1β), interleukin-6 (IL-6), and tumor necrosis factor (TNF-α), while downregulating interleukin-12p40 (IL-12p40) (Student’s t-test, *P* ≤ 0.05). We further analyzed the changes of tumor-infiltrating immune cells associated with *L. wadei*. Notably, the composition and cell numbers of myeloid cells were significantly increased (Student’s t-test, *P* ≤ 0.05) after intra-tumoral injection of *L. wadei* (Figure 5F). Among myeloid cells, granulocytic myeloid derived suppressor cells (gMDSCs) were stimulated to proliferate by the treatment of *L. wadei* (Student’s t-test, *P* ≤ 0.05). Moreover, *L. wadei* induced a pro-tumoral macrophage state. The composition and cell numbers of M2-macrophage was significantly increased, while M1-macrophage was depleted (Student’s t-test, *P* ≤ 0.05). In addition, *L. wadei* injection significantly reduced the abundance of T-bet–expressing T helper cells and regulatory T cells (Student’s t-test, *P* ≤ 0.05) (Figure 5G). T-bet has been implicated in antitumor immune responses that suppress CRC metastasis and tumor growth^35^.

To confirm that these effects on myeloid cells are specific to *L. wadei* and not a general feature of Gram-negative bacteria, we injected *P. intermedia* and *Escherichia coli MG1655* into the subcutaneous tumor model. Both *P. intermedia* and *E. coli MG1655* groups have no significant difference in tumor size with control group (Wilcoxon rank-sum test, *P* > 0.05) (Figure 5H, 5I, and Figure S4D). gMDSCs were stimulated to proliferate by the treatment of *P. intermedia* and *E. coli MG1655* (Student’s t-test, *P* ≤ 0.05) (Figure 5J). Notably, the composition and cell numbers of M1-macrophage and M2-macrophage exhibited no significant difference compared to control group after *P. intermedia* and *E. coli MG1655* treatment (Student’s t-test, *P* > 0.05) (Figure 5J).

In summary, we found that *L. wadei* and its secretome promote CRC tumorigenesis. Furthermore, *L. wadei* induces immune suppression in the tumor microenvironment and M2 macrophage polarization, whereas gMDSCs expansion appears to be a more general response to the presence of Gram-negative bacteria.

### *Leptotrichia wadei* Promotes M2 Macrophage Polarization via Metabolic Reprogramming

We next performed single-nucleus RNA sequencing (snRNA-seq) on tumors treated with *L. wadei* or Phosphate-buffered saline (PBS) to investigate potential regulatory roles of different cell types. Gene expression-based clustering revealed a clear separation between the *L. wadei* and PBS-treated tumor groups (Figure 6A). A total of 107,248 nuclei representing 22,480 genes were captured and visualized using Uniform Manifold Approximation and Projection (UMAP). Based on previously established marker genes (Figure S5A), the cells were categorized into 10 distinct clusters, comprising various epithelial, stromal, and immune cell types (Figure S5B). Given that *L. wadei* has been shown to promote CRC progression and induce M2 macrophage polarization in the *in vivo* model, we focused our further analyses on the macrophage population. A total of six macrophage subtypes were identified, including monocyte-derived cells, M1-like macrophages, M2-like macrophages, M1/M2 intermediate macrophages, cycling M2-like macrophages, and C1qa/b/c-expressing M2-like macrophages (Figure 6B). Next, RNA velocity was used to predict the future states of individual cells (Figure 6C). This analysis indicated the potential differentiation pathway on the macrophage population. Particularly, macrophages within the tumor microenvironment treated of *L. wadei* exhibited an enhanced propensity for M1-like macrophages to transition towards an M2-like state.

**FIGURE 6.**
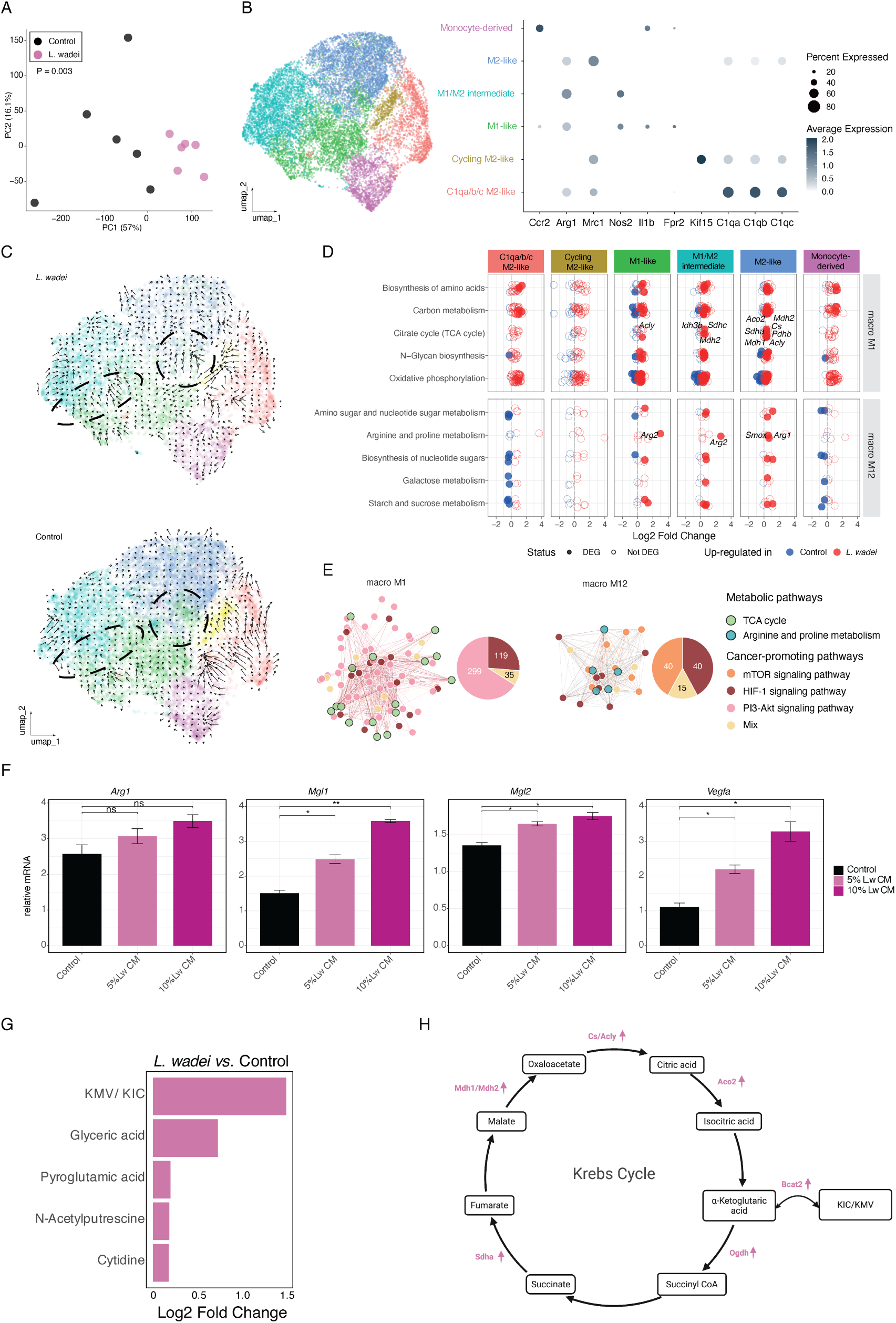
*Leptotrichia wadei* promotes M2 macrophage polarization via metabolic reprogramming. (A) Principal component analysis (PCA) of the transcriptional profiles of tumors treated by *L. wadei* and PBS (Control) from the mice. P value was calculated through the PERMANOVA. (B) UMAP plot of the macrophage population of tumors collected from the mice, colored by sub-clusters (left). Dot plots represent the identified marker genes to indicate sub-cell types (right panels). (C) RNA velocity generated with scVelo of *L. wadei* and PBS (Control) treated tumor’s macrophage populations. Arrows represent the cell trajectories across clusters, inferring differentiation cell trajectories. The two circled areas represent the transition inference M1-like macrophages to M1/M2 intermediate macrophages and M2-like macrophages respectively. (D) Modules M1 and M12 from weighted gene co-expression network analysis (WGCNA) of macrophage populations. Average log2(fold-change) profile of the genes in all enriched pathways per module for *L. wadei* versus Control. Red and blue colors indicate up- and down-regulated genes in *L. wadei* treated tumors, respectively. Significant expressed genes were exhibited by the filled nodes, while non-significant genes were shown by circles. (E) Co-expression networks based on TCA cycle and arginine and proline metabolism associated genes in module M1 and M12 respectively. Each node represents one gene, and each edge refers to the co-expression relationship between two connected nodes. Node color reflects pathway association. Colored linkages indicate the pathway origin of genes linked with tumor promoting signaling pathways. (F) mRNA expression of *Arg1*, *Mgl1*, *Mgl2*, and *Vegfα* was measured in bone marrow derived macrophages after stimulation with original culture medium (Control), 5% *L. wadei* conditioned medium (Lw CM) and 10% Lw CM, respectively for 24 hours (n=2/group). (G) Bar plot of increased metabolites in *L. wadei* conditioned medium vs. original condition medium (POMA, P ≤ 0.05, FDR ≤ 0.25, log2FC ≥ 0). Metabolites also increased in *G. morbillorum* are removed. (H) Schematic of tricarboxylic acid cycle (TCA) cycle in M2-like macrophage on snRNA data. Metabolic network was adapted from the KEGG pathways (https://www.genome.jp/kegg/). Upward arrows next to genes indicate increased expression in *L. wadei*-treated tumors compared with PBS-treated tumors. Created with Biorender.com.

To obtain a more comprehensive view of the transcriptional regulation of macrophage populations, we performed weighted gene co-expression network analysis (WGCNA), identifying 12 gene modules (Figure 6D). We then analyzed differentially expressed module genes between *L. wadei*- and PBS-treated tumors and assessed KEGG pathway enrichment for each macrophage subtype (FDR ≤ 0.05). In module M1, the top enriched KEGG pathways included biosynthesis of amino acids, carbon metabolism and the citrate cycle (TCA cycle). Genes belonging to the second half of the TCA cycle, including *Sdha*, *Mdh1*, *Mdh2*, *Acly*, *Aco2*, and *Idh3b*, were significantly upregulated in M2-like macrophages, indicating an enhanced TCA cycle in these cells in *L. wadei*-treated tumors. In module M12, arginine and proline metabolism were enriched in M2-like macrophages, with *Arg1* and *Smox* showing significantly up-regulation in *L. wadei*-treated tumors.

We further examined gene co-expression patterns within these modules (Figure 6E). In module M1, genes involved in the citrate cycle (TCA cycle) were co-expressed with genes assigned to the PI3K–Akt and HIF-1 signaling pathways. The PI3K–Akt pathway showed the highest number of connections (n = 334) and is known to contribute to tumor initiation and progression^36^. In module M12, genes involved in arginine and proline metabolism were co-expressed with genes in the mTOR and HIF-1 signaling pathways. Both mTOR and HIF-1 signaling are key cancer pathways that coordinate tumor metabolism, growth, and angiogenesis, representing attractive targets for therapy^37,38^.

As the *L. wadei* secretome was found to drive the transition of M1-like macrophages toward an M2-like macrophages *in vivo* and snRNA data, we next asked if it exerts a direct effect on M1 macrophage populations. To validate this hypothesis, we incubated M1 macrophages induced from bone marrow derived macrophage cells with *L. wadei* conditioned medium. Compared with the control group, this treatment significantly increased the expression of the M2 macrophage markers Mgl1 and Mgl2, as well as the pro-tumoral marker Vegfα, while Arg1 showed an upward trend (Figure 6F). To decipher the active constituent(s) of *L. wadei* conditioned medium, we next performed untargeted metabolomics analysis comparing the conditioned medium of *L. wadei* and *G. morbillorum* to the original culture medium (POMA, P ≤ 0.05, FDR ≤ 0.25, log2FC ≥ 0) (Figure 6G). Among the metabolites uniquely secreted by *L. wadei*, 3-Methyl-2-oxovaleric acid (KMV) and 4-Methyl-2-oxovaleric acid (KIC) were the most enriched metabolites and have been reported to induce a pro-tumoral macrophage state^39^. Cross-checking these insights with our snRNA data, we observed *Bcat2* upregulation (FDR ≤ 0.05) in M2-like macrophages after *L. wadei* treatment (Figure 6H). Given that *Bcat2* links KIC/KMV metabolism with α-ketoglutarate (αKG), this may indicate enhanced TCA activity, supported by increased expression of *Ogdh*, *Sdha*, *Mdh1*/*Mdh2*, *Cs*, *Aco2*, and *Acly*.

Overall, we observed that *L. wadei* treatment promotes the transition of M1 macrophages toward an M2-like state, alongside secretion of the branched-chain keto acids KIC and KMV by *L. wadei*. In these tumors, upregulation of the TCA cycle and arginine/proline metabolism in M2-like macrophages is linked to key tumor-promoting signaling pathways.

## DISCUSSION

In this study, we integrated multi-omics data to investigate the role of gut microbiome in 77 patients with CRC to address five key questions: (1) which stage-associated microbial and metabolic signature in stool and blood distinguish early from late CRC; (2) which functional traits characterize these stool microbial signatures and link gut microbiome variation to systemic host metabolism; (3) how tumor- and normal mucosa–resident microbial communities differ from stool microbiome in composition and functional potential; (4) whether tumor- and mucosa–enriched microbes resolve into distinct metabolic capacity that reflect tissue colonization niches; and (5) whether species-specific metabolic outputs within these niches can be experimentally linked to functional driver or passenger roles in CRC progression. By leveraging the multi-omics depth of this cohort and applying advanced sequencing and analytic methods, our study provides the breadth and resolution needed to systematically assess the contribution of the gut microbiota to CRC.

Consistent with previous studies, we observed that specific gut microbes, including *E. coli*, *B. fragilis*, *P. stoma*tis, and *P. micra* are enriched in late-stage CRC and have been validated *in vivo* using animal models^31,40–42^. Beyond these previously reported taxa, we identified additional stool-strain biomarkers: *H. porci*, *M. smithii* (reported to be enriched in metastatic CRC^13^), *CAG-170* (family *Oscillospiraceae*), and *A. celatus_A* enriched in late-stage CRC, whereas *A. rectalis*, *B. massiliensis*, and *G. qucibialis* were enriched in early-stage CRC. In the stool mycobiome, we observed a shift in diversity from stage I to later stages (II–IV), driven by enrichment of *M. restricta*, *C. albicans*, *R. pusillus*, and *R. emersonii*, consistent with prior evidence implicating *M. restricta* and *C. albicans* in CRC progression and metastasis^43,44^.

Serum metabolomics revealed continuous stage-associated changes in metabolites, including increases in dihydrothymine and acetylcholine^45,46^, and decreases in 3-indoxylsulfuric acid reflecting a systemic metabolic signature linked to CRC progression. Importantly, stool bacterial functional clusters defined using KEGG Orthology modules, displayed stage–specific metabolic focuses with late-stage-enriched clusters particularly enriched for energy metabolism pathways, including the TCA cycle. These findings underscore the interconnection between stool microbial composition, functional potential, and host systemic metabolism during CRC development^46–48^. However, some functional clusters showed no detectable associations with serum metabolites, suggesting limited measurable systemic effects. Consistently, the taxa driving these clusters were largely non–tissue-associated: 72% were not detected in tissue and 94% were not significantly different between tumor and normal mucosa, implying that their functional signatures may primarily reflect colonization niches rather than play a major role in CRC progression.

At the tissue level, the CRC tumor microenvironment harbors a multi-domain microbiome that contributes to immune evasion and CRC progression^49^. Deep shotgun metagenomic analysis revealed tumor- and normal-mucosa-specific bacterial species and functional modules distinct from stool samples. Tumor tissues were enriched for oral-associated bacteria, particularly *Streptococcus* and *Leptotrichia* species with co-occurring KEGG modules for amino acid transport (M00227, M00229, M00226, M00225) and anaerobic regulation (M00456), suggesting adaptation to nutrient-rich hypoxic tumor niches^50^. Metabolomic profiling of bacterial species enriched in the tumor and normal mucosa revealed four distinct metabolic clusters, with tumor-enriched and mucosa-enriched species largely segregating. Within the tumor-resident cluster, we observed taxa reported to promote tumorigenesis (*Klebsiella pneumoniae*) in mice models^51^, co-occurred with *Streptococcus* taxa without demonstrated tumor-promoting activity (*Streptococcus anginosus*, *Streptococcus parasanguinis*, and *Streptococcus sanguinis*)^6^, suggesting that shared high-level functional profiles may capture niche preference more than impact on CRC progression.

One mixed cluster included both tumor- and mucosa- enriched species, including *B. fragilis*, *P. micra*, and *C. butyricum* previously reported as either pro- or anti-tumorigenic^21,31,32^. Functional testing confirmed that the butyrate-producing species *A. hadrus* and *C. butyricum* suppressed CRC cell growth and survival^52^, mirroring the effects of butyrate controls. In contrast, within the same cluster, we identified *L. wadei*, a previously uncharacterized species in CRC, as a driver of tumor growth *in vivo* by inducing M2 macrophage polarization. Single-nucleus RNA sequencing further confirmed the M1-to-M2 macrophage transition, with upregulation of TCA cycle and arginine/proline metabolism in M2-like macrophages, linked to key tumor promoting signaling pathways (PI3K-Akt, HIF-1 and mTOR). Moreover, the *L. wadei* secretome alone is sufficient to drive M2 macrophage polarization *in vitro*. Further metabolomic profiling showed that the *L. wadei* secretome was uniquely enriched for the branched-chain keto acids KIC and KMV compared to *G. morbillorum*. KIC and KMV have been reported to promote a pro-tumoral macrophage state and to fuel macrophage TCA cycle activity^39^, consistent with the upregulated TCA cycle observed in the snRNA data. These results highlight that general metabolic cluster membership identifies candidates but individual species-specific outputs are necessary to pinpoint functional drivers of CRC progression.

In summary, we systematically characterized microbiome, mycobiome, metabolome, lipidome, and cytokine changes across stool, blood, and paired tumor and normal mucosa tissues to better delineate CRC progression. Strain-level analyses revealed distinct early- and late-stage stool microbial signatures, which grouped into functional clusters with characteristic metabolic outputs. Further bacterial culturing with untargeted metabolomics revealed clusters largely reflect colonization niches, distinguishing tumor- versus mucosa-enriched species. Through targeted *in vitro* and *in vivo* studies, we demonstrated that species-specific metabolic outputs, rather than global community features, define functional roles as drivers or passengers. Specifically, we identified the oral bacterium *L. wadei* as a functional driver of CRC progression, promoting tumor growth via induction of M2 macrophage polarization and upregulation of tumor-promoting metabolic and signaling pathways. Together, these results provide a framework to distinguish colonization patterns from functional contributions, revealing how species-specific metabolic properties determine the impact of individual gut microbes on CRC and offering a strategy to prioritize candidate microbial drivers for experimental validation.

### Limitations of the study

One limitation of this study is that the patient cohort was exclusively recruited from Germany. Including a multi-center, international cohort in future studies could enhance the generalizability and applicability of the findings across diverse populations.

## RESOURCE AVAILABILITY

### Lead contact

Further information and requests for resources and reagents should be directed to and will be fulfilled by the Lead Contact, Gianni Panagiotou.

### Materials availability

This study did not generate new unique reagents.

### Data and code availability

The sequencing data generated in this paper have been deposited in NCBI SRA and GEO, and are publicly available. The metabolome and lipidome data are available at the NIH Common Fund’s National Metabolomics Data Repository (NMDR, supported by NIH grant U2C-DK119886) website, the Metabolomics Workbench, https://www.metabolomicsworkbench.org/. The accession number is listed in the key resources table. New codes were not generated in this study. Additional information required to reanalyze the data reported here is available from the lead contact upon request.

## ACKNOWLEDGMENTS

The project was funded by the Federal Ministry of Research, Technology and Space (BMFTR, Germany) under the project PerMiCCion (Project ID 01KD2101A). We would like to thank for their support the Deutsche Forschungsgemeinschaft (DFG, German Research Foundation) under the project “MICROVERSE II” (EXC 2051/2 – Project-ID 390713860) and the European Union’s Horizon Training Mobility Actions (TMA)- Marie Skłodowska-Curie Action Doctoral Networks (MSCA-DN) under the grant agreement No. 101169068. The ColoCare Study was supported by National Institutes of Health (NIH)/National Cancer Institute (NCI) grant U01206110. We thank the Typas lab at EMBL for kindly providing a subset of the bacterial isolates used in this study. We are thankful to the Core Facility Functional Genomics at FLI for high throughput imaging.

## AUTHOR CONTRIBUTIONS

Conceptualization: G.P. and B.G.; methodology: L.X., B.S., Z.Z., and T.S.; biological experiments: T.K., S.T., V.D, M.M., and K.R.; investigation: L.X., B.S., Z.Z., T.S., S.T., Y.N., and V.D; visualization: L.X., B.S., and Z.Z.; data curation: L.X., B.S., Z.Z., T.S., T.K., S.T., V.D, and M.M.; writing – original draft: L.X., Z.Z., and G.P.; writing—review & editing: all co-authors; funding acquisition: G.P.; resources, G.P., B.G., C.E.Z., A.B., Y.N., C.C-M., M.Z., C.M.U.; supervision: G.P. and B.G..

## DECLARATION OF INTERESTS

The authors declare no competing interests.

## SUPPLEMENTAL INFORMATION

**FIGURE S1.**
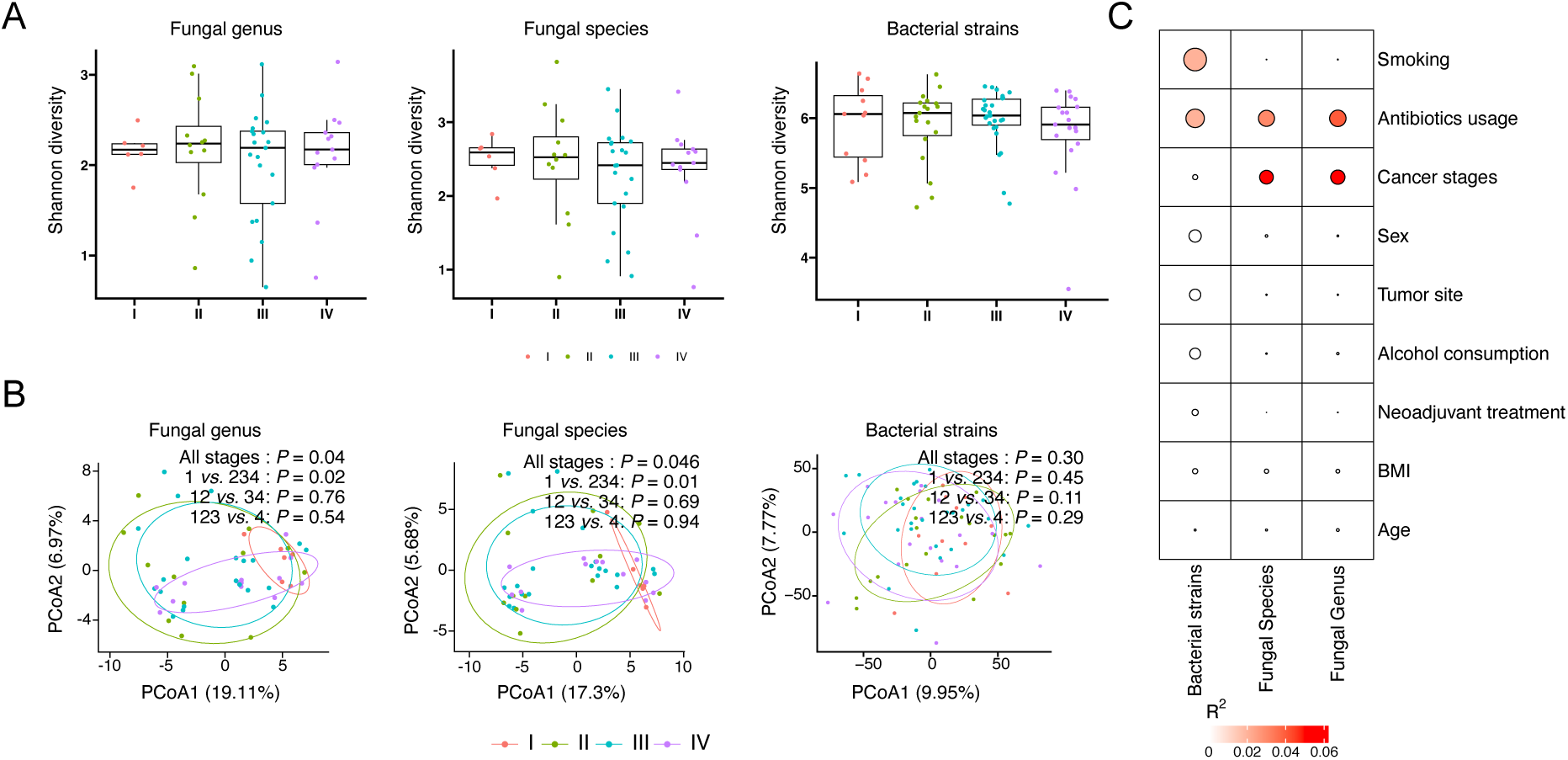
Diversity of stool fungal and bacterial community (related to Figure 1). (A) Box plots of alpha diversity (Shannon index) for fungal genus and species level and bacterial strain level were computed. Centre lines denote the median value, boxes contain the Q1 and Q3 quartiles (IQR). The whiskers extend up to 1.5 × IQR, and values beyond these bounds are considered outliers. Significance test between cancer stages was performed using two-sided Wilcoxon rank-sum test. None of the indices is significant (raw P > 0.05). (B) Principal coordinate analysis (PCoA) of beta diversity (robust Aitchison distance) for fungal genus and species level and bacterial strain level. Statistical testing was performed using PERMANOVA. Raw P values are shown. (C) Heatmap showing the association between clinical variables and microbial profiles at the fungal genus/species level and bacterial strain level. Statistical testing was performed using PERMANOVA. R^2^ describes estimated explained variance. Raw P values are shown.

**FIGURE S2.**
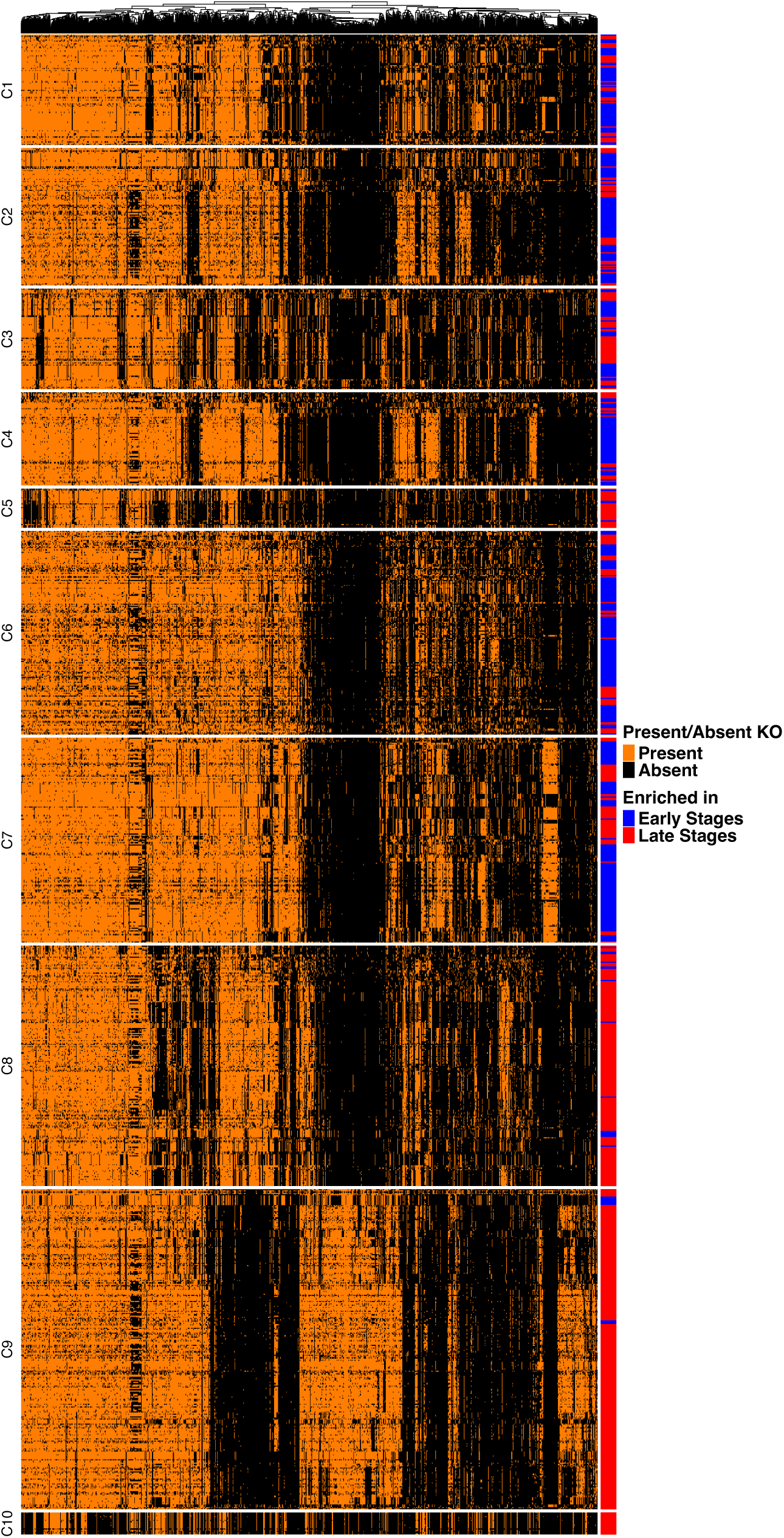
Functional cluster of signature bacterial strains (related to Figure 2). Presence/absence matrix of Kyoto Encyclopedia of Genes and Genomes (KEGG) Orthologies (KOs) identified from signature bacterial strains. The strains are clustered by Partitioning Around Medoids (PAM) algorithm.

**FIGURE S3.**
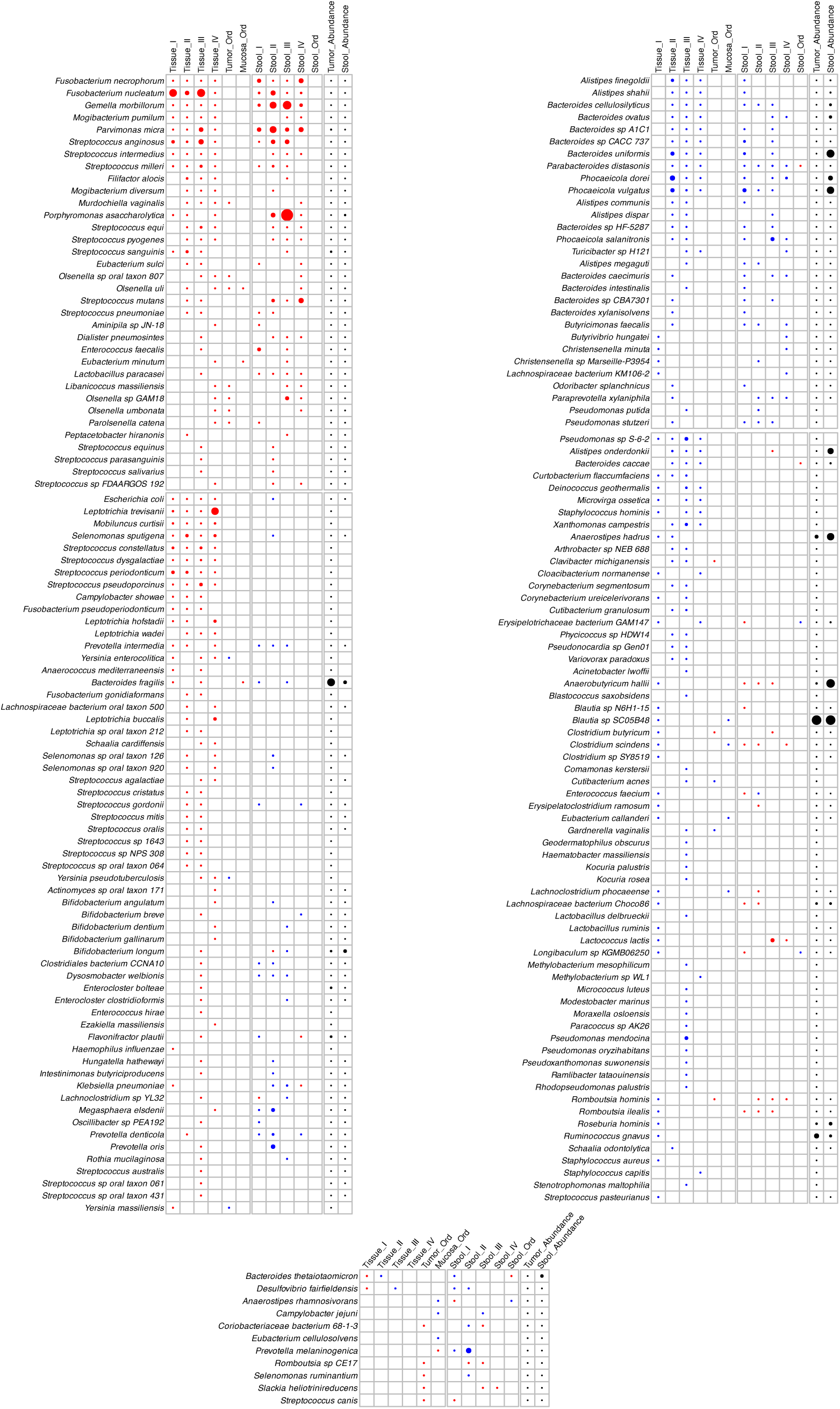
Shared and tissue-specific bacterial species in stool and tissue samples (related to Figure 3). Heatmap displaying the significantly differential bacterial signatures between tumor and normal mucosa (tissue samples), and their patterns of change in stool samples from CRC patients and controls across four cancer stages. The top two panels display bacterial signatures detected in both tissue and stool samples, while the next two panels show signatures identified exclusively in tissue samples. The bottom panel display bacterial signatures with a mixed pattern across four cancer stages in tissue samples. Circle color indicates the direction of enrichment (tumor *vs.* normal mucosa), while circle intensity corresponds to the statistical significance (−log_10_P-value). Differential bacterial signatures were identified using DESeq2 and ordinal regression. DESeq2 results are considered significant with raw P ≤ 0.05 and FDR ≤ 0.25, while ordinal regression results are significant with raw P ≤ 0.05.

**FIGURE S4.**
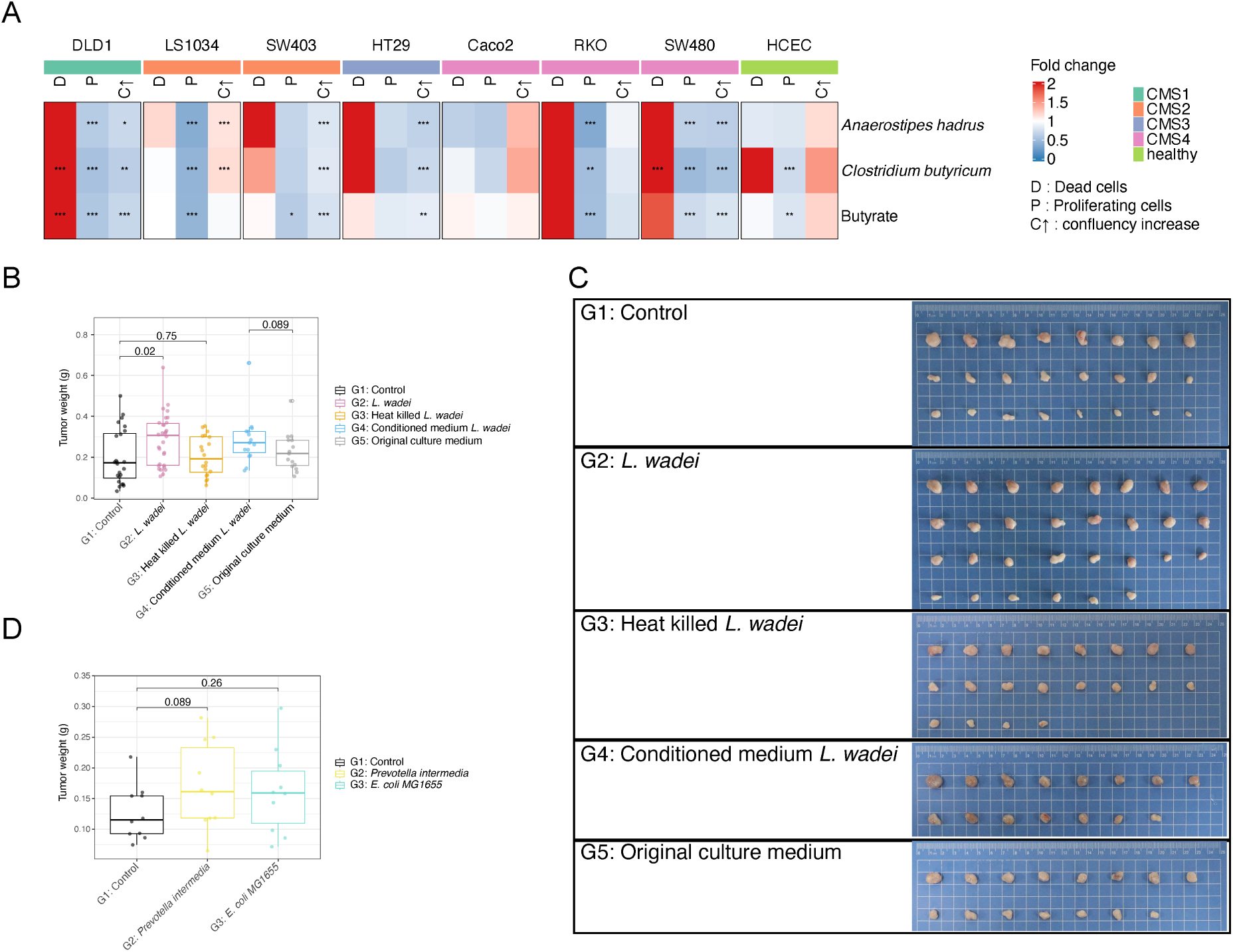
Validation of metabolic unique cluster C2 in Fig4 (related to Figure 4 and 5). (A) *In vitro* assay of *Anaerostipes hadrus* and *Clostridium butyricum* secretomes on CRC cell survival and proliferation in comparison to unspent bacterial medium control. (B) C57BL/6J mice with MC38 subcutaneous tumors were injected with Control (sterile PBS (with 0.5% cysteine), n = 24), *L. wadei* (1 x 10^6^ CFU) (n = 30), heat killed *L. wadei* (n = 20), *L. wadei* conditioned medium (n = 15), or original culture medium (n = 15) twice a week. The tumor weights of indicated groups at day 17 were compared by using Wilcoxon rank-sum test. (C) Images of representative MC38 subcutaneous tumors after the intra-tumoral injection of the indicated groups in C57BL/6J mice. (D) C57BL/6J mice with MC38 subcutaneous tumors were injected with gram negative bacteria (*Prevotella intermedia* and *E. coli MG1655*). The tumor weights of indicated groups at day 17 were compared by using Wilcoxon rank-sum test. Each group has 10 mice.

**FIGURE S5.**
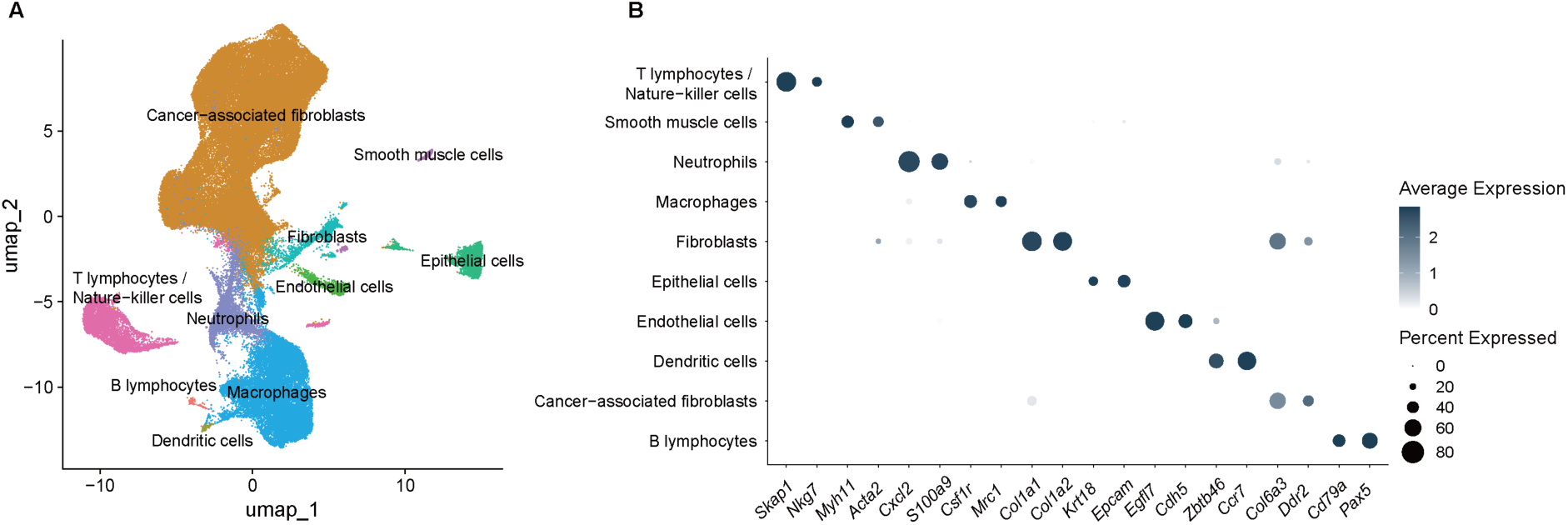
Cell type annotation based on marker genes identified through snRNA sequencing (related to Figure 6). (A) UMAP plot of the tumors collected from the mice, colored by sub-clusters. (B) Dot plots represent the identified marker genes to indicate cell types.

**Table S1.** Patient metadata.

| Table S1 Patient metadata. |  |  |  |  |  |  |
| --- | --- | --- | --- | --- | --- | --- |
|  | N | I, N= 11<sup>a</sup> | II, N= 19<sup>a</sup> | III, N= 28<sup>a</sup> | IV, N= 19<sup>a</sup> | p-value<sup>b</sup> |
| Sex | 77 |  |  |  |  | 0.5 |
| female |  | 5 / 11 (45%) | 9 / 19 (47%) | 9 / 28 (32%) | 5 / 19 (26%) |  |
| male |  | 6 / 11(55%) | 10 / 19 (53%) | 19 / 28 (68%) | 14 / 19 (74%) |  |
| Age | 77 | 67 (9) | 66 (9) | 62 (12) | 59 (11) | 0.079 |
| BMI | 77 | 28.9 (4.6) | 26.7 (5.2) | 26.8 (6.1) | 26.6 (5.2) | 0.3 |
| Tumor site | 77 |  |  |  |  | 0.085 |
| Colon |  | 7 / 11 (64%) | 8 / 19 (42%) | 6 / 28 (21%) | 8 / 19 (42%) |  |
| Rectum |  | 4 / 11 (36%) | 11 / 19 (58%) | 22 / 28 (79%) | 11 / 19 (58%) |  |
| Neoadjuvant treatment | 77 |  |  |  |  | < 0.001 |
| No |  | 10 / 11 (91%) | 19 / 19 (100%) | 19 / 28 (68%) | 7 / 19 (37%) |  |
| Yes |  | 1 / 11 (9%) | 0 / 19 (0%) | 9 / 28 (32%) | 12 / 19 (63%) |  |
| Neoadjuvant method | 22 |  |  |  |  | 0.005 |
| chemotherapy |  | 0/1 (0%) | 0 / 0 | 0 / 9 (0%) | 8 / 12 (67%) |  |
| radiochemotherapy |  | 1/1 (100%) | 0 / 0 | 9 / 9 (100%) | 4 / 12 (33%) |  |
| Alcohol consumption | 75 | 16 (23) | 20 (32) | 14 (34) | 10 (10) | 0.9 |
| Antibiotics usage | 70 |  |  |  |  | 0.056 |
| No |  | 10 /10 (100%) | 13 / 18 (72%) | 21 / 24 (88%) | 11 / 18 (61%) |  |
| Yes |  | 0 / 10 (0%) | 5 / 18 (28%) | 3 / 24 (13%) | 7 / 18 (39%) |  |
| Smoking | 75 |  |  |  |  | 0.2 |
| Current |  | 1 / 11 (9.1%) | 6 / 18 (33%) | 4 / 27 (15%) | 3 / 19 (16%) |  |
| Former |  | 5 / 11 (45%) | 8 / 18 (44%) | 6 / 27 (22%) | 7 / 19 (37%) |  |
| Never |  | 5 / 11(45%) | 4 / 18 (22%) | 17 / 27 (63%) | 9 / 19 (47%) |  |

<sup>a</sup>Mean (SD); n / N (%)
<sup>b</sup>Kruskal-Wallis rank sum test; Pearson's Chi-squared test

**Table S2.** Presence of viable *L. wadei* within the *L. wadei*-injected tumors.

| Group | Animal ID | Tumor weight (g) | CFU/mL | CFU/g |
| --- | --- | --- | --- | --- |
| G2: *L. wadei*,<br>1×10<sup>6</sup> CFU | 900 | 0,1064 | 5,28E+04 | 4,96E+05 |
|  | 914 | 0,3939 | 1,80E+05 | 4,57E+05 |
|  | 969 | 0,1420 | 5,80E+03 | 4,08E+04 |
|  | 097 | 0,1494 | 1,45E+04 | 9,71E+04 |
|  | 052 | 0,4566 | 9,70E+04 | 2,12E+05 |
|  | 079 | 0,3927 | 1,02E+05 | 2,60E+05 |

## STAR★ METHODS

### EXPERIMENTAL MODEL AND STUDY PARTICIPANT DETAILS

#### Study design

This study used biospecimens and data from the ColoCare study (ClinicalTrials.gov Identifier: NCT02328677)^23^—a multicenter, international prospective cohort, recruiting newly diagnosed CRC patients before primary surgery with the goal to investigate predictors of cancer recurrence, survival, treatment toxicities, and health-related quality of life. Patients (eligibility: newly diagnosed, age 18–80 y, stages I–IV, German speaking, and able to provide informed consent) from the ColoCare Heidelberg cohort were included, recruited between June 2016 and December 2021 at the Heidelberg University Hospital, Heidelberg, Germany. Patients were staged according to the American Joint Committee on Cancer system based on histopathologic findings. Both colon (International Classification of Diseases C18) and rectal or rectosigmoidal cancer patients (International Classification of Diseases C19 or C20) were included. ColoCare has been approved by the ethics committee of the Medical Faculty at the University of Heidelberg. All study participants provided written informed consent.

#### Biospecimen collection

Primary colon cancer tissue and matched distant non-neoplastic colon tissue (at the farthest longitudinal surgical margin, 10-15 cm apart) were collected. Samples were snap frozen in liquid nitrogen within 45 min and stored at −80 °C until further processing. In addition, paired serum and stool samples were collected. Blood was primarily taken one day before surgery. Blood samples were processed within four hours and stored at −80 °C until further processing. Stool samples were collected within a week before surgery and stored at −80 °C until further processing.

#### Mice

Female C57BL/6J mice (5 weeks old) were purchased from Jiangsu GemPharmatech Co., Ltd, China. All mice were maintained in specific-pathogen-free (SPF) conditions with approval by the Institutional Animal Care and Use Committee (IACUC) of Gempharmatech Co., Ltd (AP#:GPTAP20240708-2). Mice were between 6 and 12 weeks at time of inoculation for all experiments. The food of all animals accorded with standard diet for rodents. The animals used in study were compliant with all relevant ethical regulations regarding animal research.

#### Bacteria and culture

*Leptotrichia wadei* (DSM 19758) were purchased from BNCC (BeNa Culture Collection). The strain was introduced from DSMZ and then subcultured at BNCC. *Leptotrichia wadei* were inoculated into modified Gifu Anaerobic Medium (GAM) supplemented with clarify rumen fluid. Frozen strains will be first revived by streaking onto agar plates and incubated at 37℃ for 12-72 h. Single colony will be picked and cultured in 3 mL of liquid medium broth and incubated at 37 ℃ for 12-72 h, and cells will be collected and suspended in 60% glycerin for storage with 3 replicates. PCR and sequencing are performed for strain identification. The relationship between OD values and cell numbers is estimated before animal experiments. Briefly, 1 mL of fresh cultures will be used to measure OD value every 2-8 h, the same volume of bacterial solutions will be centrifuged, washed once, and suspended to 1 mL, spreaded onto agar plates, and incubated for colony counts. Srains will be cultured overnight in liquid medium, and *L. wadei* conditioned medium was collected when OD600 of bacteria culture medium is 1. Live *L. wadei* will be washed with PBS (with 0.5% cysteine) and cell density will be adjusted to 2 × 107 CFU/mL. To obtain heat killed *L. wadei*, live *L. wadei* will be washed with PBS (with 0.5% cysteine) and the OD600 will be adjusted to 1, followed by heating at 100°C for 30 min. *Gemella morbillorum* (DSM 20572), *Parvimonas micra* (DSM 20468), *Streptococcus anginosus* (DSM 20563), *Bacteroides fragilis* (DSM 2151), *Streptococcus sanguinis* (DSM 20567), *Klebsiella pneumoniae* (DSM 30104), *Streptococcus mitis* (DSM 12643), *Streptococcus oralis* (DSM 20627), *Fusobacterium gonidiaformans* (DSM 19810), *Streptococcus salivarius* (DSM 20560), *Streptococcus parasanguinis* (DSM 6778), *Yersinia pseudotuberculosis* (YPIII), *Olsenella uli* (DSM 7084), *Bifidobacterium dentium* (DSM 20436), *Alistipes shahii* (DSM 19121), *Bacteroides cellulosilyticus* (DSM 14838), *Bacteroides ovatus* (ATCC 8483), *Bacteroides uniformis* (DSM 6597), *Parabacteroides distasonis* (DSM 20701), *Phocaeicola dorei* (DSM 17855), *Phocaeicola vulgatus* (DSM 1447), *Bacteroides caccae* (DSM 19024), *Anaerostipes hadrus* (NT5243), *Clostridium butyricum* (NT5317), *Bacteroides caecimuris* (I48), *Fusobacterium nucleatum subsp. animalis* (ATCC 51191), *Fusobacterium nucleatum subsp. nucleatum* (ATCC 25586), and *Fusobacterium nucleatum subsp. vincentii* (ATCC 49256) were inoculated into Gut microbiota medium (GMM). Frozen glycerol stocks of gut bacteria were streaked on pre-reduced BHI-blood agar or mGAM agar and incubated at 37℃ anearobically for 48 hr. Single colony of gut bacteria was inoculated into 300 uL pre-reduced Gut microbiota medium (GMM) in a 96-well plate and incubated anearobically at 37 °C for 48 hr as preculture. Fermentation was initiated by transferring 15 uL preculture into 1.5mL GMM (1:100 dilution) in a 96-well deep plate and incubated anearobically for 24hr. Optical density was determined by a plate reader (Synergy4, BioTek) at 595 nm on 100 uL the fermentation broth. The fermented broth was centrifuged 3200 ctf for 10 minutes, and the supernatant was transferred to another 96-well deep plate and stored at -80 °C freezer before further processing for CRC cell line assays and metabolomics measurement.

#### Incubation of cells with bacterial secretomes

The colorectal cancer cell lines DLD-1, SW-403, RKO, LS1034, Caco-2, HT-29 and SW-480 as well as healthy colonic epithelial cells (HCEC) were cultured in high glucose DMEM supplemented with 10% Fetal Bovine Serum and seeded in 384 well plates with 1000 cells/well for Caco-2, DLD-1, and RKO, 2000 cells per well for SW-480 and HT-29 and 4000 cells/well for LS1034 and SW403. After 24 h of incubation at 37°C, plates were imaged using an IncuCyte SX5 in 10x magnification. Cells were treated with bacterial secretomes, diluted 1:8 with cell culture medium. Butyrate (final concentration 2 mM) as well as unspent bacterial medium were used as a control. After 72 h of incubation at 37°C, plates were imaged again using an IncuCyte SX5.

#### Double staining for dead and proliferating cells

Cells were stained with 10 μM EdU solution (Invitrogen EdU Alexa Fluor 488 C10627) for 2 h at 37°C, stained with dead cell staining dye (diluted 1:700 with PBS, Invitrogen Live/Dead fixable Far Red Dead Cell Stain Kit L10120) for 30 minutes at room temperature protected from light, fixed with 4% formaldehyde for 60 minutes at room temperature and permeabilized with 0.2% Triton X-100 in PBS for 20 minutes at room temperature. Afterwards, cells were stained with 20 μL of Click-iT® Plus reaction cocktail for 30 minutes and with 15 μL of DAPI (0.33 mg/L in PBS) for 10 minutes at room temperature protected from light. Between each staining, cells were washed twice with PBS. Afterwards, cells were imaged using an ImageXPress Micro Confocal (MD) Automated Microscope (Molecular Devices) at 10x magnification.

Confluency in IncuCyte images was analyzed using the AI Confluence Analysis module within the IncuCyte software. Cell count, number of proliferating cells and number of dead cells were analysed from ImageXPress images using an algorithm created in the MetaXpress Custom Module Editor Pipeline, developed with MetaXpress software.

#### Tumor models and treatments

Five to seven weeks female C57BL/6J SPF mice were used for the subcutaneous tumor model. C57BL/6J mice were injected with 1.2 x 10^5^ MC38 colon carcinoma cells. When tumors grew up to ∼ 50 to 100 mm^3^, mice were randomly assigned and subcutaneously (s.c.) injected with 50 uL sterile PBS (with 0.5% cysteine), heat killed *L. wadei*, conditioned medium *L. wadei*, original culture medium, or 1 x 10^8^ cells (*G. morbillorum* and *L. wadei*) and 1x10^6^ cells of bacteria (*L. wadei*, *P. intermedia*, and *E. coli MG1655*) twice a week for 3 weeks. Next, tumor volumes and dissolved tumors were recorded followed by further analysis. Tumor volume (mm^3^) was calculated by using formula 0.5 x (longest diameter) x (shortest diameter). The ethical endpoint of subcutaneous tumor volume was above 4000 mm^3^ according to previous reports.

### METHOD DETAILS

#### Metagenomics sequencing of human CRC tissue and stool samples

DNA from tumor and normal mucosa tissues of CRC patients was extracted using the Ultra-Deep Microbiome kit following the manufacturer’s recommendations (Molzym, Germany) with minor modifications: initial proteinase K digestion was extended from 10 to 20 minutes, centrifugation steps were at 15000 xg, and final elution was done in 40 uL of the provided deionized water heated to 70 °C. Note that total provided tissue sample was used in each case, and negative controls used the kit-included TSB solution as input. Gel electrophoresis using Tapestation 2200 (Genomic DNA and D1000 screentapes, Agilent, USA) was used to validate product size and purity of a subset of DNA extracts. DNA concentration was measured using Qubit dsDNA HS Assay kit (Thermo Fisher Scientific, USA). Extracted DNA was fragmented to approximately 550 bp using a Covaris M220 with microTUBE AFA Fiber screw tubes and the settings: Duty Factor 10 %, Peak/Displayed Power 75 W, Cycles/burst 200, Duration 40 s and Temperature 20 °C. The fragmented DNA was used for metagenome preparation using the NEB Next Ultra II DNA library preparation kit (New England Biolabs, USA). The DNA library was paired-end sequenced (2 x 150 bp) on a NovaSeq S4 system (Illumina, USA). DNA from stool samples was extracted using the TGuide S96 Magnetic Soil/Stool DNA Kit (Tiangen Biotech (Beijing) Co., Ltd.) according to manufacturer instructions. The DNA concentration of the samples was measured with the Qubit dsDNA HS Assay Kit and Qubit 4.0 Fluorometer (Invitrogen, Thermo Fisher Scientific, Oregon, USA). The library was constructed with the VAHTS® Universal Plus DNA Library Prep Kit for Illumina according to manufacturer instructions. Paired-end sequencing of the library was on an Illumina Novaseq6000 (2×150 bp).

#### Bacterial taxonomic and functional profiling

Read quality control was performed using sunbeam (v2.1.0)^53^, including removal of low-quality reads (Phred score > 15 over 4 nucleotides), adaptors, and decontamination. To remove human-host related reads, Sunbeam mapped reads using bwa mem (v0.7.17)^54^ against the human reference genome GRCh38. For tissue samples, we further removed human-host related reads that mapped against the human pan-genome T2T-CHM13v2.0 (bowtie2 v2.5.1^55^ --very-sensitive). Tissue samples yielded an average of 181 million raw reads per sample; after host-read removal, about 4.7 million non-host reads per sample remained for downstream analyses. Bacterial taxonomic profiling was performed using KrakenUniq (v1.0.2)^56^ with default parameters. For bacterial functional annotation of stool and tissue samples, Shogun (v1.0.8)^57^ aligned reads against the RefSeq database version 82 at 95% similarity threshold. Reads mapped to multiple references were resolved to the last common ancestor (LCA) using the burst aligner. The resulting functional abundance profiles were further normalized to the median read-depth for downstream analysis using Shogun normalize.

#### Human cancer signature annotation

We followed the GATK best practices for somatic short variant detection with matched normal samples. Raw reads were trimmed using Trimmomatic v. 0.39^58^ to remove adapters and low-quality segments (LEADING:5 TRAILING:5 SLIDINGWINDOW:5:25 MINLEN:60). Reads shorter than 60bp were discarded. Reads were aligned to human reference genome (GRCh38) using bwa mem v. 0.7.17^54^. Duplicated reads were marked using Picard Tools MarkDuplicates v. 2.27.5. To minimize the impact of bacteria reads, deduplicated human-aligned reads with high-quality alignment against a comprehensive microbial pan-genome (ChocoPhlAn v.31 database from HUMAnN 3.7) were removed. In the following, we used GATK v. 4.4.0^59^. To create a reference of expected mutations (panel of normals), we started with GATK BaseRecalibrator using known SNP sites from the GATK hg38 bundle (dbsnp, 1000G, and hapmap), and used the result to apply recalibration to human-read alignments. The final panel of normal was created using GATK Mutect2 on only mucosa mapping files and --max-mnp-distance 0 followed by CreateSomaticPanelOfNormals with known allele frequencies (af-only-gnomad). Somatic variant candidates were called in matched tumor-normal mode of GATK Mutect2, including the previously generated PON and known allele frequencies. We then estimated Germline contamination rates in tumor samples using GATK GetPileupSummaries and CalculateContamination, leveraging “small_exact_common_3”. GATK FilterMutectCalls identified high-confidence somatic mutations for downstream analysis. Mutational signatures were estimated in R using signature refitting from the MutationalPatterns package (v3.8.1)^60^ and COSMIC v. 3.2 signatures. Only samples with confident signatures refit (cosine similarity >= 0.9) were kept. Each of the resulting SBS signatures was tested for differences in signature contribution across cancer stages using Kruskal-Wallis-Test.

#### Metagenome binning and quality assessment

Assembly of metagenomic reads per sample or run was performed using megahit v.1.1.3^61^ with the default parameters. After assembly, metaWRAP v.1.3.2^62^ was used to bin the assemblies using the binning module with the options ‘--maxbin2 --metabat2 --concoct’. metaWRAP was then also used for Bin refinement and individually re-assembling each bin, using the ‘bin_refinement’ module and “reassemble_bins” module respectively, with the options ‘-c 50 -x 10’, which set the quality thresholds at ≥50% completeness and ≤10% contamination. The MAGs from metaWRAP bin refinement were dereplicated at a species level using dRep v.3.4.2^63^ with dRep dereplicate and the options ‘-pa 0.9’ (primary cluster at 90%), ‘-sa 0.95’ (secondary cluster at 95%), ‘-nc 0.1’, ‘-comp 50’ (completeness threshold of 50%) and ‘-con 10’ (contamination threshold of 10%). Taxonomic classification of each MAG and isolate was performed using the GTDB-Tk v.2.1.1^64^ classify workflow with ‘gtdbtk classify_wf’ using the GTDB database release 207, which spans 317,542 genomes grouped into 65,703 species clusters. Coverage was estimated using CoverM v.0.6.1^65^ and the options ‘--min-read-percent-identity 0.99’, ‘--min-read-aligned-percent 0.75’, ‘--proper-pairs-only’, ‘--min-covered-fraction 0’ and ‘--exclude-supplementary’. MAGs were first annotated using Prokka v.1.14.6^66^ to predict protein-coding sequences. We then used eggNOG v.2.1.11^67^ to functionally characterize the MAGs using the option ‘--evalue 0.001’. A phylogenetic tree was constructed using Phylophlan v.3.1.68^68^ using its database of 400 universal amino acid marker sequences, and the ‘–diversity high’ option. ‘phylophlan_write_config_file’ was run with the ‘-d n –db_aa diamond –map_dna blastn –map_aa diamond –msa muscle –trim trimal –tree1 iqtree –tree2 raxml --force_nucleotides’ options, and the final phylophlan run was executed using ‘phylophlan’ with the options ‘-d phylophlan –diversity high –accurate’. The resulting best-scoring RAxML tree file were imported into iTOL v.7^69^ for visualization.

#### Full length ITS sequencing of human stool samples

The concentration of extracted genomic DNA was determined by Qubit 2.0 and DNA quality was checked using gel electrophoresis. PCR reactions used 200 ng of DNA with primer sets specific to hypervariable regions ITS1F (5′-CTTGGTCATTTAGAGGAAGTAA-3′) and ITS4 (5′-TCCTCCGCTTATTGATATGC-3′). Primer sets had unique barcodes. PCR products were separated on gels, and fragments with the proper amplification size were extracted and purified. Purified PCR product was used as a template for library preparation. PCR products were pooled in equal amounts and then end polished, A-tailed, and ligated with adapters. Fragments were filtered with beads. The libraries were analyzed for size distribution and quantified using real-time PCR. Sequencing of the library was performed on the PacBio sequel II.

#### Fungal full length ITS annotation

SMRTlink (v11) was applied to extract CCS reads from subreads (minPasses≥5). The CCS reads obtained were demultiplexed to samples with LIMA v1.7.0 software (Pacific Biosciences). The bioinformatic analysis was conducted using the open-source pipeline NextITS v0.6.0^70^ (its_region: full; clustering_method: vsearch; otu_id: 0.98). This pipeline is designed for full-length ITS sequencing with PacBio and available for public use and can be accessed on the GitHub platform (https://Next-ITS.github.io/). Sequences were clustered into OTUs at 98% sequence similarity with VSEARCH v2.27.0^71^ and subsequently curated with LULU v.1.0.0^72^ to remove erroneous molecular OTUs. Taxonomic annotation of OTUs was obtained using the BLASTn method with BLAST v. 2.14.0+^73^ (evalue < e^-10^) against the UNITE database v10 (https://unite.ut.ee/) to represent roughly species-level entities. Taxonomic assignments were performed at 70%, 75%, 80%, 85%, 90% and 97% sequence similarity to roughly match phylum, class, order, family, genus and species level, respectively. Taxa with sequence similarity < 70% to any taxon or match e-value > e^-50^ were considered unidentified at the kingdom level.

#### Nucleus isolation from frozen samples for snRNA sequencing

100-200 mg samples were flash frozen immediately after collection for downstream processing. Tissue samples were cut into pieces < 0.5 cm and homogenized using a glass Dounce Tissue Grinder (Sigma, cat. no. D8938). The tissue was homogenized 25 times with pestle A and 5-10 times with pestle B in 2 mL of ice-cold nuclei EZ lysis buffer. The sample was then incubated on ice for 5 min, with an additional 3 mL of cold EZ lysis buffer added. Then the sample was filtered through a cell strainer to remove large tissue particles. Nuclei were centrifuged at 500 g for 5 min at 4°C, washed with 5 mL ice-cold EZ lysis buffer and incubated on ice for 5 min. After centrifugation, the nucleus pellet was washed with 5 mL nuclei suspension buffer (NSB; consisting of 1×PBS, 0.04% BSA and 0.1% RNase inhibitor (Clontech, cat. no. 2313A)). Isolated nuclei were resuspended in 2 mL NSB, filtered through a 35 μm cell strainer (Corning-Falcon, cat. no. 352235) and counted. A final concentration of 1,000 nuclei per μL was used for loading on the channel.

#### snRNA-seq library preparation and sequencing

We prepared snRNA-Seq libraries with Chromium Next GEM Single Cell 3ʹ Reagent Kits v3.1 on the Chromium Controller (10× Genomics). We prepared single nucleus suspensions from cultured cell lines and nuclei were suspended in PBS containing 0.04% BSA. The nuclei suspension was loaded onto the Chromium Next GEM Chip G and ran the Chromium Controller to generate single-cell gel beads in the emulsion (GEMs) according to the manufacturer’s recommendation. Captured nuclei were lysed and the released RNA was barcoded through reverse transcription in individual GEMs. Barcoded, full-length cDNA was generated and libraries were constructed according to the performer’s protocol. The quality of libraries was assessed by Qubit 4.0 and the Agilent 2100. Sequencing was performed on the Illumina NovaSeq X with a sequencing depth of at least 50,000 reads per nucleus and 150 bp (PE150) paired-end reads (performed by Biomarker Technologies Corporation, Beijing, China).

#### Human serum and bacteria supernatant sample preparation for LC-MS

Sample analysis was carried out by MS-Omics as follows. For semi-polar metabolite analysis, serum samples (50 μL) were diluted with a mixture of acetonitrile, methanol, formic acid and stable isotope labelled internal standards (200 μL 1.00:0.99:0.01 v/ v/ v). The extracts were then passed through a phosphor lipid removal cartridge (Phree, Phenomenex) using centrifugation (1 400 rpm, 4°C, 10 min). An aliquot (100 μL) was transferred into a high recovery HPLC vial and the solvent was removed under a gentle flow of nitrogen. Extracts were reconstituted in a reverse phase mobile phase mixture (100 μL, 10% B in 90% A). For lipidomics analysis, serum samples (10 μL) were transferred to a Spin-X® filter (0,22 µm, Costar) and diluted in a mixture of isopropanol, SPLASH® Lipidomix® (Avanti Polar Lipids) and butylated hydroxytoluene (90 μl, 96:4 vol/vol + 10 µg). The extract was then left at room temperature for 10 min before storage at -20°C overnight. Samples were brought to room temperature and filtered by centrifugation (14 000 rpm, 5°C, 2 min). An aliquot (25 μL) was transferred into a high recovery HPLC vial and an equal parts mixture of mobile phase eluent A and B (75 µL) was added. For quality control, a mixed pooled sample (QC sample) is created by taking a small aliquot from each sample. This sample is analyzed with regular intervals throughout the sequence. For bacteria supernatant samples, 5µL sample were diluted in 115 mobile phase eluent A (10mM ammonium formate + 0.1% formic acid in ultrapure water) fortified with stable isotope labelled standards before analysis.

#### LC-MS method of human serum and bacteria supernatant samples

Sample analysis was carried out by MS-Omics as follows. For semi-polar metabolites, the analysis was carried out using a Thermo Scientific Vanquish LC coupled to a Q Exactive™ HF Hybrid Quadrupole-Orbitrap, Thermo Fisher Scientific. For bacteria supernatant samples, the mass spectrometer instrument is Orbitrap Exploris 240 MS, Thermo Fisher Scientific. An electrospray ionization interface was used as ionization source. Analysis was performed in positive and negative ionization mode. The UHPLC was performed using an adapted version of the protocol described by Doneanu et al^74^. Metabolomics processing was performed untargeted using Compound Discoverer 3.0 (Thermo Scientific) for peak picking and feature grouping, followed by an in-house annotation and curation pipeline written in MatLab (2021b, MathWorks). Identification of compounds was performed at four levels; Level 1: identification by retention times (compared against in-house authentic standards), accurate mass (with an accepted deviation of 3ppm), and MS/MS spectra, Level 2a: identification by retention times (compared against in-house authentic standards), accurate mass (with an accepted deviation of 3ppm). Level 2b: identification by accurate mass (with an accepted deviation of 3ppm), and MS/MS spectra, Level 3: identification by accurate mass alone (with an accepted deviation of 3ppm). For lipid analyses from human serum, lipidomics high-performance liquid chromatography (HPLC), coupled with high-resolution mass spectrometry (HRMS) analysis was performed using a Thermo Scientific Vanquish LC coupled to Thermo Q Exactive™ HF Hybrid Quadrupole-Orbitrap mass spectrometer (Thermo Scientific, RRID: SCR_020545). The chromatographic separation of lipids was carried out on a Waters® ACQUITY Charged Surface Hybrid (CSH™) C18 column (2.1 x 100 mm, 1.7 µm). The column was thermostated at 55°C. The mobile phases consisted of (A) Acetonitrile/water (60:40) and (B) Isopropanol/acetonitrile (90:10), both with 10 mM ammonium formate and 0.1% formic acid. Lipids were eluted in a two steps gradient by increasing B in A from 40 to 99% over 18 min. The flow rate was kept at 0.4 mL/min. Ionization was performed in positive and negative ionization mode using a heated electrospray ionization interface. The mass spectrometer operated at a resolution of 120,000 in a scan range of 200 to 1500 m/z. Iterative data-dependent MS/MS (dd-MS²) acquisition was achieved on a TopN of 10 with stepped collision energy (NCE) of 20, 40, and 60. Peak areas were extracted using Compound Discoverer 3.0 (Thermo Scientific). Compound annotations were performed against the mzCloud (Thermo Scientific) MSMS library, the Human Metabolome Database 4.0, and the MS-Omics lipid library covering 17 classes (CerP, DG, DGDG, Fatty acids, LPC, LPE, MG, PA, PC, PE, PG, PI, Plasmenyl-PC, Plasmenyl-PE, PS, SM and TG). Identification of compounds was performed at four levels; Level 1: identification by retention times (compared against in-house authentic standards), accurate mass (with an accepted deviation of 3 ppm), and MS/MS spectra, Level 2a: identification by retention times (compared against in-house authentic standards and lipid class behavior), accurate mass (with an accepted deviation of 3ppm). Level 2b: identification by accurate mass (with an accepted deviation of 3 ppm), and MS/MS spectra, Level 3: identification by accurate mass alone (with an accepted deviation of 3 ppm). Lipid annotations were curated manually by comparing retention time behavior over the number of tail chain carbons and double bonds as annotated.

#### Cytokine profiling

The concentrations of cytokines in human serum were measured by BioLegend LEGENDplex kits (LEGENDplex™ HU Essential Immune Response Panel, LEGENDplex™ HU Immune Checkpoint Panel 1 - S/P, LEGENDplex™ HU Proinflam. Chemokine Panel 1, LEGENDplex™ HU Proinflam. Chemokine Panel 2, LEGENDplex™ Human Growth Factor Panel) according to the manufacturer’s instructions. The samples were run on a Beckman Coulter Cytoflex (3L, 5V-5B-3R) and analyzed using Biolegend software (legendplex.qognit.com). The concentrations of cytokines in mouse tumors were measured by cytokine bead array (BCA). Tumor tissues were lysed using T-PER (Thermo, 78510) to obtain protein supernatants. Total protein concentration was determined by the BCA method. Concentrations of 13 cytokines, including CXCL1 (KC), TGF-β1 (Free Active), IL-18, IL-23, CCL22 (MDC), IL-10, IL-12p70, IL-6, TNF-α, G-CSF, CCL17 (TARC), IL-12p40, and IL-1β, were detected using the LEGENDplex™ Mouse Macrophage/Microglia Panel (13-plex) (Biolegend, 740846) kit. Experiments were performed according to the manufacturer’s instructions, and samples were detected on a Cytek NL-CLC flow cytometer and analyzed using Biolegend software (legendplex.qognit.com).

#### Flow cytometry

Colon tumors from mice were cut into small piece, then digested with Tumor Dissociation Kit (RWD, DHTE-5001) use single cell suspension preparation apparatus (RWD, DSC-400). Cell suspension was homogenized through a 70μm cell strainer. Cells were washed with FACS buffer. Then stained with FC blocker (Anti-CD16/CD32 antibody) at 4℃ for 10min. Cell surface markers (mCD45, CD3, mCD4, mCD8, mCD335, mCD25, mCD11b, MCHII, mF4/80, mCD206, mCD11c, mLy6G, mLy6C) were stained on ice for 60 min. Next, cells were fixed and permeablized by Intracellular Fixation & Permeablization buffer set (eBioscience, 00-5523-00), then intracellular markers (FOXP3, Ki67, GZMB, T-bet and RORgt) were stained at 4°C for 30 min. Wash twice with perm buffer, then resuspended the cells with 300 µL facs buffer, cell samples were performed on flow cytometer (Cytek Northern Lights-CLC, Cytek) and Flowjo (V10) was used to analyze the flow cytometry data.

### QUANTIFICATION AND STATISTICAL ANALYSIS

#### Decontamination of tissue microbiome

OTU table from tissue samples at species level was filtered to remove microbial species with less than 0.01% relative abundance, leaving 323 unique species out of 6,009 original species found across all samples. Species table were subsequently stringently decontaminated using the decontam package (v1.20.0)^75^ in R. Putative contaminants were identified using both final library DNA concentrations (collected independently for every sample and technical replicate) and negative blank samples (that is, ‘‘method=combined’’ in decontam; P* = 0.5); in this manner, taxa were labeled as putative contaminants for either being more abundant in negative blanks than in biological samples or for having a strong negative correlation between read fraction and DNA concentration across many biological samples. The main two assumptions of this decontamination framework are (i) that contaminants are consistently added across samples (e.g., from technician handling or reagents) and (ii) that contaminants are overall lowly abundant than authentic microbial constituents. After decontamination, 304 species were available for downstream analyses across all samples.

#### Diversity analysis

Diversity calculations were performed using the R package vegan v.2.6-6.1^76^. The alpha diversity indices of bacterial and fungal communities were calculated using Shannon index. For beta diversity, analyses were performed using the robust Aitchison distance to overcome compositionally related biases. Principal Coordinate Analysis (PCoA) were used to visualize the bacterial and fungal profile. Between-group differences were evaluated using PERMANOVA (adonis2 from vegan) with 999 permutations.

#### Procrustes Analysis

To assess overall correspondence across different omics, we performed Procrustes analysis using the R package vegan v.2.6-6.1^76^. We used robust Aitchison’s distance on microbial taxonomic data and Euclidean distance on metabolomics and lipidomics data as input to the Procrustes analysis. The significance of rotation agreement was obtained using the ‘protest()’ function with 999 permutations.

#### Clustering Analysis

Clustering analysis was performed using the ConsensusClusterPlus package (v1.64.0)^77^. The Partitioning Around Medoids (PAM) algorithm was applied, with 80% of the genes randomly resampled over 1000 iterations. The number of potential clusters (K) ranged from 2 to 10. The optimal number of clusters was determined using the calcICL function, selecting the maximum K where all clusters exhibited a consensus score greater than 0.8. For distance calculation, binary distance was used for the microbial functional profile, while Pearson distance was applied for the metabolomics profile. Microbial functional clusters were classified as “pure” when >70% of strains within a cluster were enriched in either early-stage or late-stage strains. Overrepresentation of KEGG pathways in these pure functional clusters was tested using clusterProfiler (v4.8.3)^78^ with pvalueCutoff = 0.05 and qvalueCutoff = 0.05. KOs were considered present in a cluster if they occurred in ≥50% of strains in that cluster. Finally, only KEGG pathways uniquely enriched in either early-stage–enriched or late-stage–enriched pure clusters were retained.

#### Untargeted metabolomics/lipidomics data normalization

In both samples from human cohort and bacterial culture supernatants, we filtered out metabolites and lipids with Descriptive Power (DP) ≤ 2.5, and replaced value < LOD (The Limit of Detection) with NA. Missing values in metabolome and lipidome profiles were imputed by LOD/2. In the human cohort, we filtered out metabolites and lipids with missing percentage > 30% in all samples before missing value imputation. For human cohort, metabolite and lipid data within annotation level 1, level 2a and level 2b were used as confidently annotated compounds. For bacterial culture supernatants, metabolite data within annotation level 1 and level 2a were used as confidently annotated compounds. The bacterial supernatant samples were normalized to their corresponding original culture medium. Specifically, normalization was performed by calculating the log2-transformed ratio of the mean abundance from the bacterial supernatant to the mean abundance from the culture medium. All data were logarithmically transformed before further analysis.

#### Metabolite Enrichment Analysis

Confidently annotated compounds, defined as those with annotation level 1 or 2a, were used to identify secreted or consumed metabolites for each bacterial strain. Significantly secreted or consumed metabolites were determined using the POMA package (v 1.10.0)^79^ in R (P ≤ 0.05, FDR ≤ 0.25). Metabolites that were secreted or consumed by more than 50% of the strains within a cluster were considered common secreted or consumed metabolites. The overrepresentation of KEGG pathways associated with these common metabolites was assessed using MetaboAnalyst 6.0^80^.

#### snRNA-Seq data processing

We performed alignment to this amended reference using 10x Cell Ranger v7.2, which employs the STAR sequence aligner. The reference genome was the human genome GRCm39 or the mouse genome mm39. We determined gene expression counts using unique molecular identifiers (UMIs) for each cell barcode-gene combination. Following alignment, we filtered cell barcodes to identify those which contain nuclei using the approach implemented in Cell Ranger, and only these barcodes were considered for downstream analysis. To remove the nuclei with low quality, we applied seurat (v5.1.0)^81^ R package to construct 4 objects according to the four tissues and filter nuclei with gene number over 200 and below 5,000, as well as the ratio of mitochondria lower than 5% were maintained, and genes with at least one feature count in more than ten nuclei were used for the following analysis. Doublets were identified performing the DoubletFinder (v2.0.4)^82^ R package with principal components (PCs) 1-20. nExp was set to 0.08 × nCells2/10,000 and pN to 0.25. pK was determined through paramSweep_v3, and cells classified as doublets were removed before downstream analysis. In total, 22,480 genes were identified in 107,248 cells of 12 samples. We then adopted the Seurat-implemented NormalizeData function to perform the library-size correction and logarithm transformation, with the obtained expression matrix used for downstream analyses. We then adapted the workflow of Seurat to perform dimension reduction and unsupervised clustering. First, the top 2000 highly variable genes (HVGs) were selected by the FindVariableFeatures function (Seurat) with the parameter selection.method = “vst”. Effects of the total UMI count and mitochondrial gene percentage were then regressed out from the HVG expression matrix with the ScaleData function (Seurat). Subsequently, data were integrated and removed batch effects using CCAIntegration method, and the Principal Component Analysis was then performed by the RunPCA function (Seurat) on the scaled HVG expression matrix, retaining the top 30 components for downstream analyses.

#### Cell type annotation, cell sub-clustering and differentially expressed gene (DEG) identification

FindMarkers function (avg_log2FC ≥ 0.25 and p_val_adj < 0.05) was used to find significantly expressed markers of each cell cluster. The information of marker genes for overall cell types was shown in **Supplementary figure S5**. subset function was applied to extract macrophage cells, which was then analyzed through the same data processing workflow previously completed for the overall cells. Final macrophage sub-clustering data including 15,271 cells for the downstream analysis. As for the DEG identification in different phenotypes, Seurat’s FindMarkers function (avg_log2FC ≥ 0.25 and p_val_adj < 0.05) was used to identify the significantly different expressed genes between the *L. wadei* and control treatment groups at the cellular levels.

#### RNA-velocity analysis and gene co-expression network construction

To limit the batch effects and the quality difference of the samples, we performed the RNA-velocity analysis for individual samples as suggested^83^. Briefly, velocyto software (v0.17.17)^84^ was conducted to recount the spliced and un-spliced unique molecular identifiers, and velocyto.R R package (v0.6) was used for RNA velocity estimation. The estimation was restricted to genes which were previously applied to data integration.

Metabolic gene co-expression networks of the macrophages were constructed using hdWGCNA (v0.3.3)^85^. Genes with 5% prevalence across cells were used for network construction. Pseudobulk metacells aggregating nuclei were created that belonged to the same sample and cell type setting a nearest-neighbour parameter k of 30 and using 10 as the maximum number of shared nuclei between metacells. Modules were identified through the ConstructNetwork function. Cytoscape (v3.10.1)^86^ with edge-weighted spring-embedded layout to visualize the networks based on the topoligcal overlap matrices (TOMs) generated by GetTOM function in hdWGCNA.

#### Statistical Analysis

R (version 4.3.0) was utilized for data analysis. Untargeted metabolomics/lipidomics data and snRNA-Seq data were logarithm transformed before downstream analyses. For bacterial and fungal taxonomic profiles, only features that appeared in more than three samples and reached a relative abundance greater than 0.01 % in at least one group were retained for downstream differential-abundance analyses. Pairwise comparisons were conducted to evaluate the progression of colorectal cancer (CRC) across consecutive stages (I *vs.* II-IV, I-II *vs.* III-IV, and I-III *vs.* IV). Bacterial and fungal taxonomic profiles were analyzed using the DESeq2 package v1.40.2^87^ in R, with fitType set to “local” and sfType set to “poscounts”. For bacterial functional profiles, library size normalization was first performed using the GMPR package (v0.1.3)^88^ in R, followed by differential abundance analysis using DESeq2 (v1.40.2)^87^ with the same parameter settings. Metabolomics and lipidomics data were analyzed using the POMA package v.1.10.0^79^ in R with default parameters. In addition, ordinal regression analysis was performed using the ordinal package v.2024.12-13^89^ in R to identify features showing progressive increases or decreases across CRC stages I to IV. This analysis was applied to bacterial and fungal taxonomic profiles, bacterial functional profiles, human cancer signatures, as well as metabolomics and lipidomics datasets. Univariate correlations involving continuous and categorical data were performed using the rank-based Spearman correlation by R package ggpubr (v0.6.0). The values were presented as the means ± SDs. The data from the two unpaired groups were compared via a two-tailed unpaired Student’s t-test. Values of *p < 0.05, **p < 0.01, and ***p < 0.001 were considered as statistically significant. The Benjamini-Hochberg procedure was applied to calculate the FDR to adjust P values for multiple hypothesis testing.

## Notes

### Competing Interest Statement

The authors have declared no competing interest.

